# Environmental filtering and habitat (mis)matching of riverine invertebrate metacommunities

**DOI:** 10.1101/2022.04.09.487317

**Authors:** David Murray-Stoker, Kelly M. Murray-Stoker, Fan Peng Kong, Fathima Amanat

## Abstract

**Aim:** Metacommunities are assembled through a combination of local and regional processes, with the relative importance of the drivers of assembly depending on ecological context. Global change can alter community assembly at both local and regional levels, potentially shifting communities into disequilibrium with their local environmental conditions. In this study, we evaluated the spatial variation in environmental filtering and habitat matching of 1078 riverine macroinvertebrate communities distributed across nine ecoregions within the conterminous United States.

**Location:** Conterminous United States.

**Taxon:** Freshwater macroinvertebrates.

**Methods:** We first quantified spatial patterns in environmental filtering, habitat matching, and functional trait diversity. We then used boosted regression trees to identify (1) functional trait predictors of environmental filtering and habitat matching and (2) environmental, landscape, and network variables that predict functional trait abundances.

**Results:** Our results demonstrated that environmental filtering but not habitat matching varied strongly by ecoregion. We also found that functional trait diversity varied by ecoregion, but not as strongly as the signatures of environmental filtering. We did not identify consistent functional trait predictors for both environmental filtering and habitat matching, with trait predictors instead varying by individual traits, trait categories, and ecoregions. Notwithstanding inconsistent trait predictors, environmental filtering was primarily influenced by habitat preference traits while habitat matching was primarily influenced by both habitat preference and dispersal traits. Predictors of functional traits also varied by trait category and ecoregion, with habitat preference and dispersal traits primarily influenced by network variables.

**Main conclusions:** Our study demonstrates the contingent patterns and drivers of environmental filtering and habitat matching on a macroecological scale. We aim for this work to provide the foundation on which trait-environment relationships can be further quantified and causal explanations established in the context of community disequilibrium and applied to conservation and management of freshwater systems.

## Introduction

Understanding and predicting community responses to anthropogenic global change is increasingly important to ecology and conservation. Communities are assembled through a combination of local (e.g., environmental conditions, habitat suitability, species interactions) and regional (e.g., dispersal, connectivity) processes (Belyea & Lancaster, 1999; Leibold et al., 2004; Leibold & Chase, 2017; Vellend, 2010), and both types of processes act in concert to drive community assembly, composition, and diversity (Cornell & Harrison, 2014; Mittelbach & Schemske, 2015; Vellend, 2010). The relative influence of local and regional processes can depend on both ecological context (Belyea & Lancaster, 1999; Leibold et al., 2004; Leibold & Chase, 2017) and on the composition and diversity of the regional species pool (Cornell & Harrison, 2014; Mittelbach & Schemske, 2015). Moreover, global change can alter the underlying drivers of community assembly and shift communities into mismatch or disequilibrium with their local environmental conditions (Blonder et al., 2015). As communities are the ultimate link between biodiversity and ecosystem functioning (Hooper et al., 2012; Isbell et al., 2017), a primary focus on the structure of and responses by ecological communities could help to understand patterns of biodiversity and identify potential ecosystem-level alterations under anthropogenic global change.

A useful approach to evaluate community responses to global change could be to focus on species niches and environmental conditions (Blonder et al., 2015; Carscadden et al., 2020). Under this community disequilibrium framework (Blonder et al., 2015), species niches and environmental conditions are used to determine the degree of environmental filtering and habitat matching. Environmental filtering occurs when local communities are composed of species with similar niches relative to the regional species pool, while environmental permissiveness is the converse: local communities composed of species with dissimilar niches compared to the regional species pool (Blonder et al., 2015; Knight et al., 2020). Habitat matching occurs when species comprising the local community have niches similar to the observed environmental conditions relative to species in the regional pool, and habitat mismatch occurs when species in the local community have niches dissimilar to the observed environmental conditions of the local community (Blonder et al., 2015; Knight et al., 2020). Environmental filtering is a common driver of community assembly (Cottenie, 2005; Grönroos et al., 2013; Heino et al., 2017; Murray□Stoker & Murray□Stoker, 2020; Soininen, 2014), whereby environmental variables constrain community structure by sorting species based on niche-based responses to biotic and abiotic factors (Belyea & Lancaster, 1999; Leibold et al., 2004; Vellend, 2010). Additionally, by determining if community composition reflects the local environmental conditions, it is possible to identify if communities are out of balance in response to environmental change (i.e., habitat mismatch; Blonder et al. 2015; Knight et al. 2020). Environmental filtering and habitat matching therefore provide complementary information regarding community assembly and disequilibrium, respectively.

Quantifying and comparing environmental filtering and habitat matching can indicate patterns of these metrics, but determining the underlying mechanisms would require further investigation. A focus on functional traits could provide a mechanistic understanding of environmental filtering and habitat matching. Functional traits characterize organismal performance in relation to habitat and resource use and links those autecologies to community and ecosystem processes (McGill et al., 2006; Spasojevic et al., 2014). For example, tolerance allows species to inhabit more disturbed (e.g., urbanized or polluted habitats: Barnum et al. 2017; Gianuca et al. 2018) or extreme (e.g., mean or range of temperature: Spasojevic et al. 2014; stream flow intermittence: Bogan et al. 2015; Schriever et al. 2015) environments. Additionally, dispersal traits can allow species to better reach isolated habitats (Germain et al., 2017; Horváth et al., 2019) or track changing environmental conditions (Grönroos et al., 2013; Heino & Grönroos, 2014). Within the context of the disequilibrium framework, a functional trait-based approach could be used to identify which traits are related to environmental filtering and habitat matching and then establish how functional traits relate to environmental conditions (i.e., trait-environment relationships sensu Legendre et al. 1997). In doing so, drivers underlying spatial patterns in environmental filtering and habitat matching could be identified to understand current trends and predict community responses under global change.

Rivers and streams are useful model ecosystems because the dendritic and hierarchical structure of these systems can affect the relative influence of local and regional processes on communities (Altermatt, 2013; Heino, Melo, et al., 2015). In rivers and streams, local drivers typically relate to water quality and physical habitat (Astorga et al., 2011; Grönroos et al., 2013; Heino et al., 2003, 2017). Regional drivers are generally related to dispersal and connectivity between habitats (Astorga et al., 2011; Heino et al., 2003, 2017; Mykrä et al., 2007; Sarremejane et al., 2017). Previous studies have frequently found that local drivers of community assembly and structure typically have a greater influence than regional drivers (Heino et al., 2003; Kärnä et al., 2015), although hierarchical assembly can be modulated by an interaction between local and regional factors (Astorga et al., 2011; Göthe et al., 2017; Heino et al., 2017; Murray Stoker & Murray Stoker, 2020). At macroecological scales, differences among ecoregions and the composition and diversity of species pools could further contextualize the identity and importance of local and regional drivers on community assembly and structure (Astorga et al., 2011; Heino et al., 2017; Murray Stoker & Murray Stoker, 2020). Importantly, rivers and streams comprise a fraction of water (0.01%) and land (0.8%) cover, but these ecosystems are habitat to a disproportionate amount of biodiversity (~6% of all described species; Dudgeon et al. 2006); however, these ecosystems and concomitant biodiversity are increasingly under threat from anthropogenic forces (Dudgeon, 2019). Specifically: overexploitation, pollution, flow modification, habitat degradation or loss, introduced species, and climate change (Booth et al., 2016; Dudgeon, 2019; Dudgeon et al., 2006). It is therefore critical to understand the patterns and drivers of community responses to global change in freshwater ecosystems.

Our objective was to identify predictors of disequilibrium-trait and trait-environment relationships that could help inform basic and applied research in riverine ecosystems. Here, we combined the disequilibrium framework with trait-environment relationships to evaluate the patterns and drivers of environmental filtering and habitat matching in riverine ecosystems at the macroecological scale. We focused on riverine invertebrate communities from 1078 sites nested within nine ecoregions of the conterminous United States. In this study, we asked four questions:

**Q1** = How do environmental filtering and habitat matching vary across ecoregions?
**Q2** = How does functional trait diversity vary across ecoregions?
**Q3** = Which functional traits are linked to environmental filtering and habitat matching?
**Q4** = What are the environmental predictors of functional trait abundances?

We used a multi-step approach to answer these questions (Appendix Figure S1). First, we quantified and compared environmental filtering and habitat matching across ecoregions (**Q1**). Second, we compared patterns of functional trait diversity across ecoregions (**Q2**). Third, we related functional traits to environmental filtering and habitat matching to identify which traits were related to disequilibrium metrics (**Q3**). Finally, we related environmental, geographic, and network predictors to functional traits to identify the drivers of functional trait structure (**Q4**).

## Methods

### Data source

We used the publicly-available dataset from the 2008-2009 National Rivers and Streams Assessment (NRSA) conducted by the United States Environmental Protection Agency (USEPA 2016a)The objective of the NRSA was to assess the biotic and abiotic condition of river and stream ecosystems (USEPA 2016a). During the NRSA, a total of 1924 sites were sampled from nine ecoregions across the conterminous United States (USEPA 2016a): coastal plains, Northern Appalachians, northern plains, Southern Appalachians, southern plains, temperate plains, western mountains, upper midwest, and xeric. Following the sampling design, the NRSA collected data on benthic macroinvertebrates, water chemistry, and physical habitat characteristics. Landscape attributes for the watershed of each site were derived from the National Hydrography Dataset Plus (USEPA 2016a, b). Due to variable selection and use of macroinvertebrate community data described below, not all data were available for all sites and the number of sites included in the analyses was reduced from 1924 to 1078 (Figure 1). Number of sites from each ecoregion included in the analyses: coastal plains, n = 100; Northern Appalachians, n = 97; northern plains, n = 134; Southern Appalachians, n = 180; southern plains, n = 125; temperate plains, n = 110; upper midwest, n = 86; western mountains, n = 134; xeric, n = 112.

**Figure 1:**
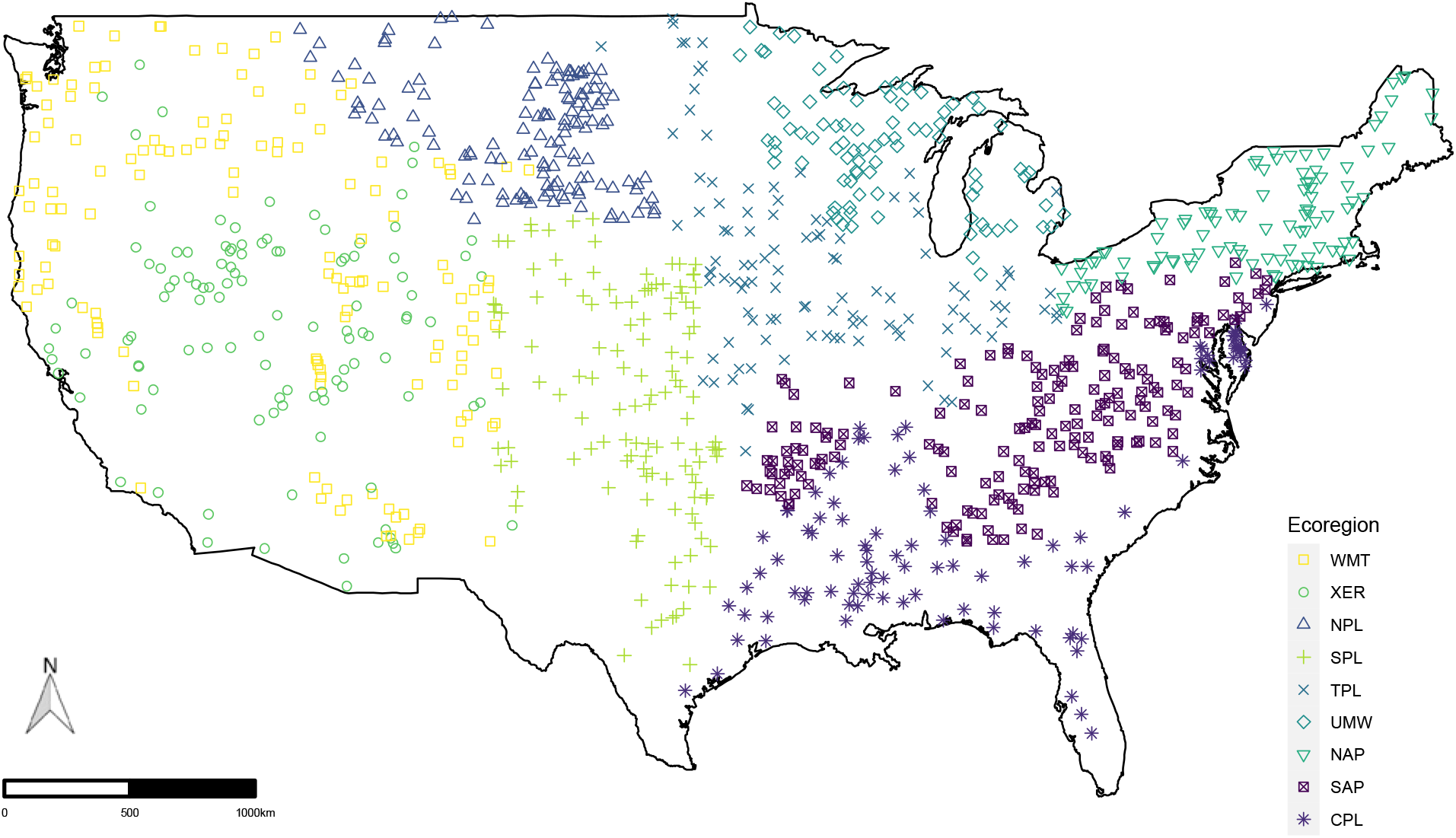
Site map of all sampling locations included in the metacommunity structure and assembly analyses differentiated by ecoregion assignment. Ecoregions are abbreviated in the legend as: coastal plains = CPL (n = 100), northern Appalachians = NAP (n = 97), northern plains = NPL (n = 134), southern Appalachians = SAP (n = 180), southern plains = SPL (n = 125), temperate plains = TPL (n = 110), upper midwest = UMW (n = 86), western mountains = WMT (n = 134), and xeric = XER (n = 112).

### Q1: How do environmental filtering and habitat matching vary across ecoregions?

#### Disequilibrium analytical framework

We used the disequilibrium analytical framework (Blonder et al. 2015) to calculate environmental filtering (*δ*) and habitat matching *λ*) metrics. In this approach, observed local communities and associated environmental conditions are compared to null expectations derived from a resampled regional pool. Species’ distributions and associated environmental conditions in the regional pool set the expected range of environmental conditions (i.e., niche) for a given species. Both *δ* and habitat *λ* are calculated by using a null model to compare how species in the local community match the observed environmental conditions relative to predictions within multidimensional niche space (Appendix Figure S1; Blonder et al., 2015).

Prior to calculating *δ* and *λ*, two statistics are quantified: (1) Δ, the observed community niche volume is measured as the niche space occupied by the community; it accounts for the niche breadth of each species in the community (Appendix Figure S1; Blonder et al., 2015). (2) Λ, the vector between the niche centroid of the observed community and the expected niche centroid of a null community (Appendix Figure S1; Blonder et al., 2015). Null distributions of Δ and Λ are generated by sampling communities of equal richness from the regional pool.

Standardized deviations (*δ* and *λ*) are determined by comparing observed Δ and Λ to expected values derived from the resampled regional pool through z-transformations:

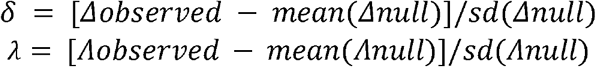

Interpretations of environmental filtering (*δ*) and habitat matching (*λ*) values are provided in Table 1.

**Table 1:** Interpretation of environmental filtering (*δ*) and habitat matching (Λ) and the ecological processes underlying the statistics.

| Statistic | Interpretation | Ecological Processes |
| --- | --- | --- |
| $\delta$ | Filtering ($\lambda < 0$) | Filtering could result from species sorting, antagonistic species interaction, or sufficient dispersal for species to reach and occupy suitable habitat |
| | Permissiveness ($\lambda > 0$) | Permissiveness could result from weak species sorting, facilitative or weak species interactions, weak dispersal that reduces species' abilities to reach and occupy suitable habitat, or extensive dispersal (i.e., mass effects) that allows species to occupy an otherwise unsuitable habitat |
| $\lambda$ | Matching ($\delta < 0$) | Matching could result from species tracking environmental changes, species interactions (antagonistic or facilitative) that cluster species in niche space, or sufficient dispersal for species to reach and occupy suitable habitat |
| | Mismatch ($\delta > 0$) | Mismatch could result from priority effects (i.e., lag between current environmental conditions and community composition), species interactions (antagonistic or facilitative) that do not cluster species in niche space, or weak dispersal that reduces species' abilities to reach and occupy suitable habitat |

#### Disequilibrium pipeline

We used the disequilibrium analytical framework to quantify environmental filtering (*δ*) and habitat matching (*λ*) for each individual community across each of the nine ecoregions. For each ecoregion, the analysis was conducted in four steps by defining: (1) the local community, (2) the regional species pool, (3) niche variables, and (4) observed environmental conditions. We defined the local community as the taxa present at a site and the regional species pool was all of the present taxa in the ecoregion. We selected four environmental variables to estimate species’ niches: (1) air T_max_ annual, (2) air T_min_ annual, (3) pH, and (4) conductivity. Temperature variables relate to the physiology and thermal niche of organisms, while pH and conductivity broadly relate to tolerance to pollution and disturbance (Dodds & Whiles, 2010; Merritt et al., 2019; Paul & Meyer, 2001; Walsh et al., 2005). A detailed description and rationale for variable selection is provided in Appendix S1. The disequilibrium analysis is sensitive to the number of niche variables selected, so we followed the guidelines suggested by Blonder et al. (2015) to include as few and ecologically- and physiologically-relevant variables as possible. Observed environmental conditions were the observed values for T_max_ annual, T_min_ annual, pH, and conductivity at each local community.

Filtering and habitat matching statistics for each local community were quantified from the mean of 100 random samples of the niche of each present taxon. To calculate *δ* and Λ, we generated 100 null communities by resampling with replacement from the regional pool while maintaining species richness. Null communities were then used to generate distributions for Δ and Λ, from which *δ* and *λ* were quantified. We conducted the above analyses using the ‘comclim’ package (Blonder, 2018), and the analysis was parallelized using the ‘snow’ package (Tierney et al., 2018). Environmental variables were centered and scaled (i.e., mean = 0, standard deviation = 1) to meet assumptions of the disequilibrium analysis.

#### Environmental filtering and habitat matching

We compared environmental filtering and habitat matching across ecoregions using one-way ANOVAs with Type III sums-of-squares, with post-hoc Tukey’s HSD tests to identify differences among groups; effect sizes calculated as eta-squared (η^2^). Vector components of habitat matching and mismatch were extracted from the disequilibrium pipeline for each of the four variables used to estimate species’ niches (i.e., T_max_ annual, T_min_ annual, pH, and conductivity). These vector components indicate which niche axes match or mismatch community composition (Blonder et al., 2015). We compared vector components by ecoregion to determine if there was spatial variation in axes of habitat matching or mismatch. Vector components were compared by ecoregion using a one-way ANOVA with Type III sums-of-squares followed by post-hoc Tukey’s HSD tests; effect sizes were calculated as η^2^.

### Q2: How does functional trait diversity vary across ecoregions?

#### Functional traits

We selected three broad functional trait categories (dispersal, habitat, and ecology) and 21 modalities related to community assembly and habitat suitability within the context of the disequilibrium framework (Appendix Table S1). We compiled a macroinvertebrate trait database using information from available databases and primary literature (Poff et al. 2006; Appendix Table S1). We used functional traits for two focal purposes: (1) identifying functional trait predictors of environmental filtering and habitat matching (i.e., Disequilibrium-by-Traits) and (2) determining environmental predictors of functional trait abundances (i.e., Trait-by-Environment); both of these approaches are described in greater detail below.

#### Functional trait diversity

We measured and compared functional trait diversity to understand the background pool of functional trait diversity. We measured functional trait diversity for each community using four indices (Laliberté & Legendre, 2010; Villéger et al., 2008): (1) functional richness (FRic); (2) functional divergence (FDiv); (3) functional evenness (FEve); and (4) functional dispersion (FDis). A detailed description of each functional trait diversity metric is provided in Appendix S1. We compared FRic, FEve, FDiv, and FDis by ecoregion using one-way ANOVAs with Type III sums-of-squares followed by post-hoc Tukey’s HSD tests; effect sizes in the ANOVAs were calculated as η^2^. We also analyzed how the effects of functional trait diversity covary with ecoregion to affect environmental filtering and habitat matching using ANCOVAs. Four separate ANCOVAs were fitted for each response variable (i.e., environmental filtering and habitat matching). Influence of predictors was estimated with Type III sums-of-squares and effect sizes were calculated as partial eta-squared (η^2^_P_).

### Q3: Which functional traits are linked to environmental filtering and habitat matching?

#### Disequilibrium-by-Traits analyses

We used boosted regression trees (BRTs) to identify which functional traits were the best predictors of environmental filtering and habitat matching. We fitted one set of BRTs with environmental filtering as the response and one set of BRTs with habitat matching as the response; functional trait abundances were fitted as the predictor variables for both Disequilibrium-by-Traits BRTs. Our goal was to identify which functional traits were useful predictors of filtering and niche mismatch, with separate BRTs for each ecoregion to allow for the relative influence of functional traits to vary by ecoregion. Disequilibrium-by-Traits BRTs were: (1) fitted to a Gaussian error distribution, (2) fitted to 10,000 trees, (3) had a learning rate of 0.0001, (4) a minimum of 5 observations per terminal node, (5) had an interaction depth of 4, (6) had a bagging fraction of 50%, and (7) used ten-fold cross validation. We fitted all BRTs using the ‘gbm()’ function in the ‘gbm’ package (Greenwell et al., 2020), with code parallelized using the ‘ snow’ package (Tierney et al., 2018). To help interpret results, we summed the relative influence by trait category; categorical sums were calculated by summing all predictor variables within the category with a relative influence greater than 5.00 (e.g., Murray-Stoker & Murray-Stoker, 2020). We used a conservative threshold to suggest evidence for an effect: predictors with relative influence > 5.00 suggest importance while relative influence < 5.00 does not suggest importance.

### Q4: What are the environmental predictors of functional trait abundances?

#### Environmental, landscape, and network variables

We selected environmental, landscape, and network variables to identify influential drivers of Trait-by-Environment relationships. We selected seven environmental variables [total nitrogen (μg L^-1^), total phosphorus (ug L^-1^), dissolved organic carbon (mg L^-1^), natural streambed cover (proportion of areal cover), large woody debris (m^3^ per 100 m stream reach), algal cover (proportion of areal cover), and macrophyte cover (proportion of areal cover)], four landscape variables [forested cover (% of basin area), agricultural cover (% of basin area), urban cover (% of basin area), and impervious surface cover (% of basin area)], and seven network variables [latitude, longitude, basin area (km^2^), mean annual flow (m^3^ s^-1^), mean basin elevation (m), range of basin elevation (m), and site centrality (distance to centroid in km)]. For a more detailed description of the environmental, landscape, and network variables, please see Appendix S1. Note: we intentionally did not use T_max_ annual, T_min_ annual, pH, and conductivity as predictors in the Trait-by-Environment relationships because those variables were used to define species’ niches in the disequilibrium pipeline. We were interested in comparing Disequilibrium-by-Traits and Trait-by-Environment relationships, and it would be inappropriate to introduce predictors in one analysis (Trait-by-Environment) that were used to define the response variables in a separate analysis (Disequilibrium-by-Traits).

#### Trait-by-Environment analyses

We used boosted regression trees (BRTs) to identify which environmental variables were the best predictors of trait abundances. We fitted separate sets of BRTs for each of the 21 functional trait states (Appendix Table S1), with each set of BRTs fitted with the trait as the response and environmental, landscape, and network (Appendix Table S2) variables as the predictors in the Trait-by-Environment BRTs. Our goal was to identify which variables were useful predictors of trait abundance to better understand how (Trait-by-Environment) and why (Disequilibrium-by-Traits) environmental filtering and habitat matching vary. Trait-by-Environment BRTs were fitted using identical parameters to the Disequilibrium-by-Trait BRTs: the only difference between the Disequilibrium-by-Traits BRTs and the Trait-by-Environment BRTs is the latter were fitted to a Poisson distribution instead of a Gaussian distribution because the response variables were counts and not normally distributed. We fitted all BRTs using the ‘gbm()’ function in the ‘gbm’ package (Greenwell et al., 2020), with code parallelized using the ‘snow’ package (Tierney et al., 2018). To help interpret results, we again summed the relative influence by predictor category; categorical sums were calculated and variable importance interpreted as described above for the Disequilibrium-by-Trait BRTs. Additionally, for the Trait-by-Environment BRTs, we counted the number of trait-by-ecoregion combinations of each predictor category to illustrate trends by environmental, landscape, and network predictors of functional traits.

All above analyses were performed using R (version 4.1.1; R Core Team 2021) in the RStudio environment (version 2021.09.0; RStudio Team 2021). All ANOVAs and ANCOVAs were evaluated using the ‘Anova()’ function in the ‘car’ package (Fox & Weisberg, 2018), and post-hoc Tukey’s HSD tests were calculated using ‘HSD.test()’ in the ‘agricolae’ package (de Mendiburu, 2021). We calculated η^2^ and η^2^_P_ using the ‘eta_squared()’ function in the ‘effectsize’ package (Ben-Shachar et al., 2020). Model assumptions were assessed using the ‘check_model()’ function in the ‘performance’ package (Lüdecke et al., 2021). Data management and figure creation were facilitated using the ‘tidyverse’ (Wickham et al., 2019) and ‘ggpubr’ (Kassambara, 2020) packages. A diagram of all analyses and how they relate to each of our research questions is provided in Appendix Figure S1.

## Results

### Spatial variation of environmental filtering and habitat matching

Environmental filtering varied strongly by ecoregion (F_8, 1069_ = 1517.496, P < 0.001, η^2^ = 0.919, Figure 2). Environmental filtering (lower *δ*) was strongest in the northern plains ecoregion, followed by the Southern Appalachians and upper midwest ecoregions, while environmental permissiveness (higher *δ*) was highest in the western mountains ecoregion followed by the Northern Appalachians. Habitat matching also varied by ecoregion, but this effect was weak (F_8, 1069_ = 2.277, P = 0.020, η^2^ = 0.017, Figure 2); post-hoc Tukey’s HSD did not identify any influential pairwise differences between ecoregions. Qualitatively, habitat mismatch (higher Λ) was stronger in the western mountains, southern plains, and northern plains ecoregions while habitat matching (lower *λ*) was stronger in the upper midwest and xeric ecoregions.

**Figure 2:**
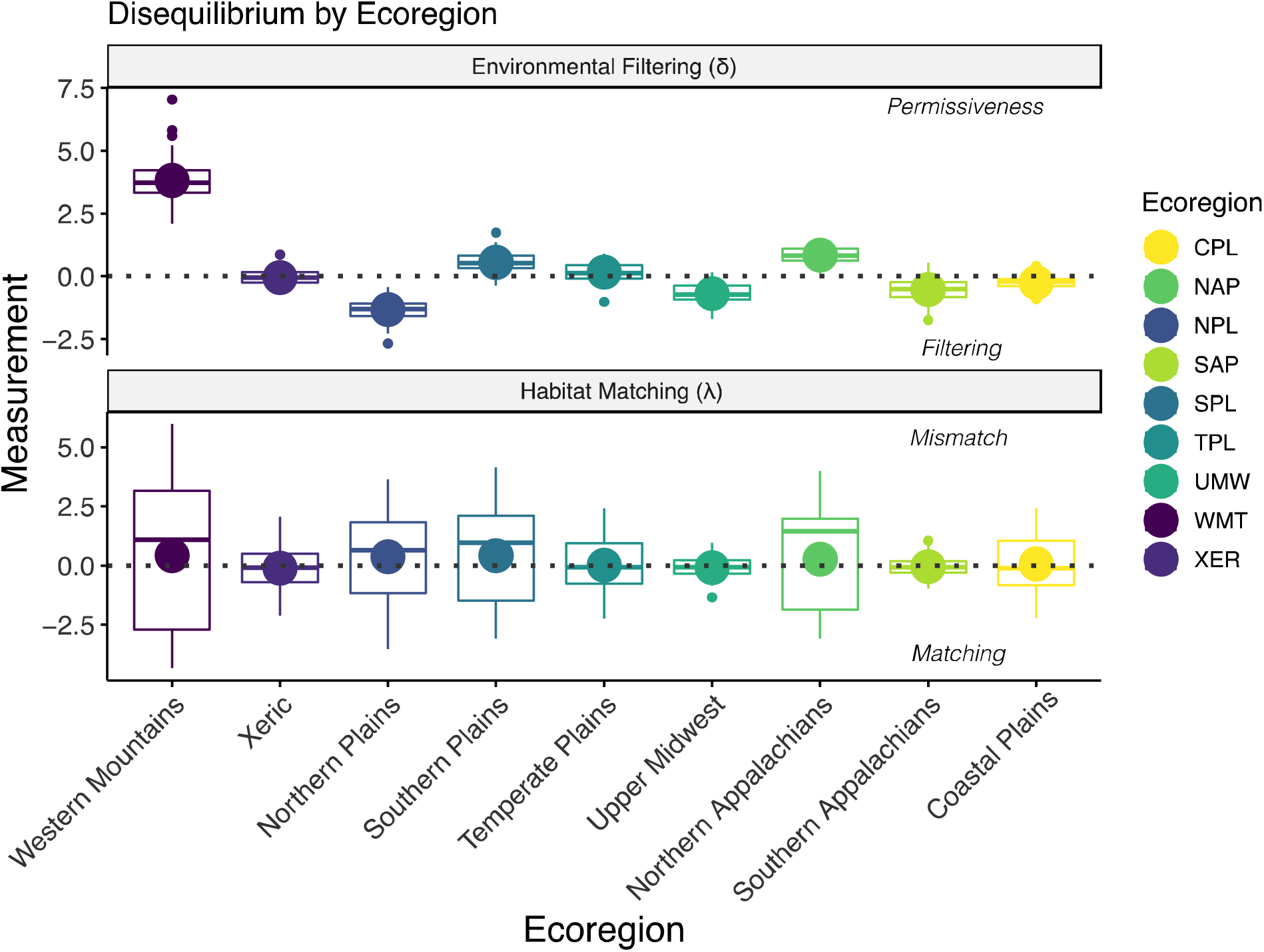
Facet plot of environmental filtering (*δ*) and habitat matching (λ) by ecoregion. Large circles represent the mean and boxplots display the interquartile range (0.25, 0.75), minimum, and maximum, with smaller circles indicating outliers. The grey dashed line indicates a value of 0. Ecoregions are arranged on the x-axis in approximate position based on longitude.

### Vectors of habitat matching and mismatch

Vector components varied by ecoregion in response to habitat matching and mismatch (Appendix Figure S2). The T_max_ annual vector components varied by ecoregion (F_8, 1069_ = 2.897, P = 0.003, η^2^ = 0.021), but post-hoc tests did not identify any pairwise differences. Generally, T_max_ annual pushed the coastal plains and Northern Appalachians ecoregions into mismatch and the xeric, southern plains, and northern plains ecoregions into habitat matching. The T_min_ annual vector components also varied by ecoregion (F_8, 1069_ = 3.555, P < 0.001, η^2^ = 0.026), driving mismatch in the temperate plains and habitat matching in the northern plains. Additionally, the pH vector components varied by ecoregion (F_8, 1069_ = 3.602, P < 0.001, η^2^ = 0.026), driving mismatch in the western mountains and habitat matching in the northern plains. There was no evidence that conductivity vector components varied by ecoregion (F_8, 1069_ = 0.920, P = 0.499, η^2^ = 0.007)

### Functional trait diversity

All four measures of functional trait diversity varied by ecoregion. Functional richness was highest in the Southern Appalachians and lowest in the northern plains, southern plains, and xeric ecoregions (F_8, 1069_ = 14.621, P < 0.001, η^2^ = 0.103, Figure 3), while FEve was highest in the southern plains and lowest in the Northern Appalachians, upper midwest, and xeric ecoregions (F_8, 1069_ = 8.089, P < 0.001, η^2^ = 0.059, Figure 3). Functional divergence was highest in the coastal plains and lowest in the xeric ecoregions (F_8, 1069_ = 6.667, P < 0.001, η^2^ = 0.049, Figure 3), and FDis was highest in the Southern Appalachians and lowest in the xeric ecoregions (F8, 1069 = 8.286, P < 0.001, η^2^ = 0.059, Figure 3).

**Figure 3:**
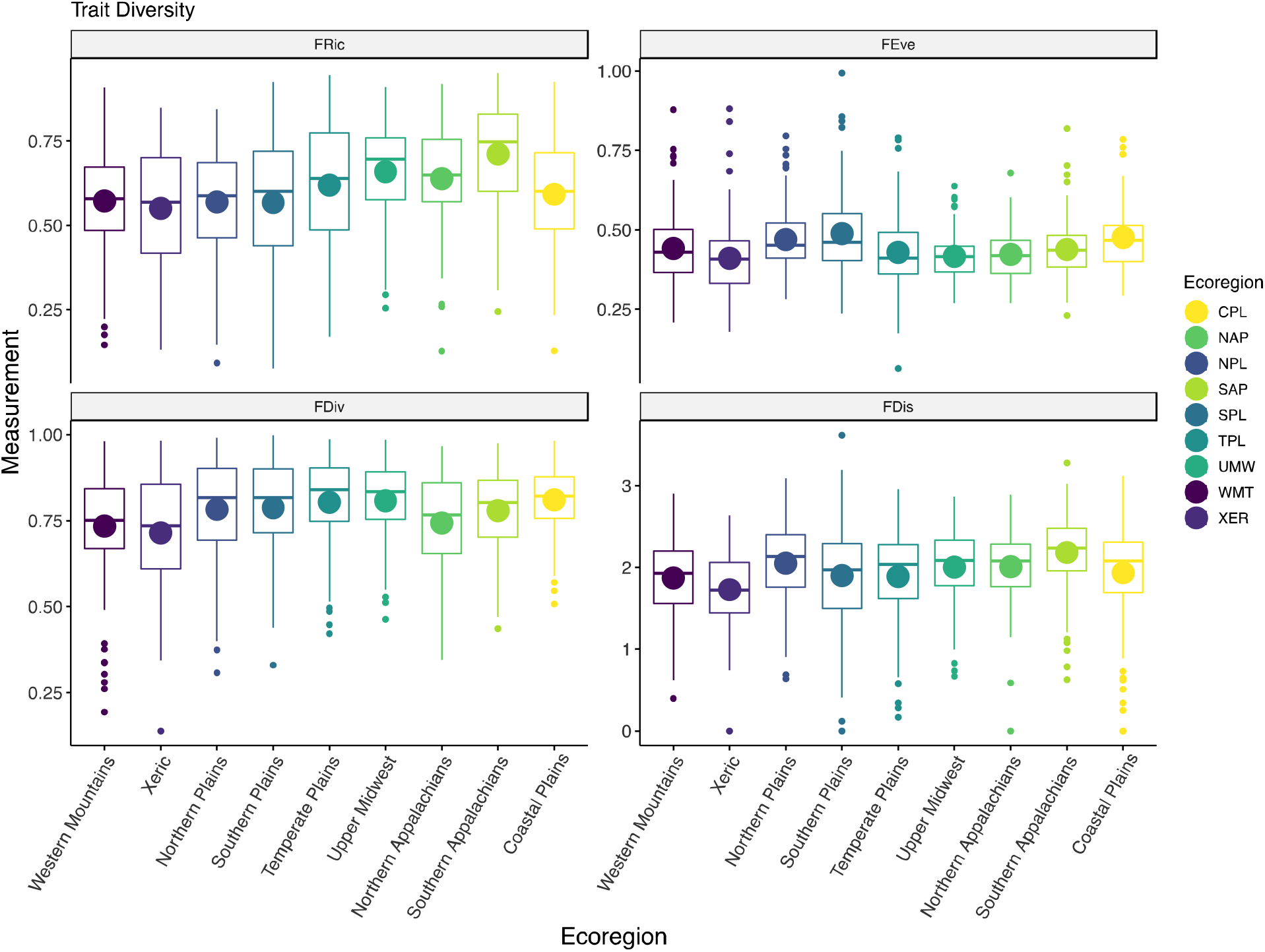
Facet plot of functional richness (FRic), functional evenness (FEve), functional divergence (FDiv) and functional dispersion (FDis) by ecoregion. Large circles represent the mean and boxplots display the interquartile range (0.25, 0.75), with smaller circles indicating outliers. Ecoregions are arranged on the x-axis in approximate position based on longitude.

### Disequilibrium and functional trait diversity

Functional trait diversity was more frequently related to habitat matching than environmental filtering (Appendix Figures S2 and S3). Environmental filtering was only predicted by the interaction of FDiv and ecoregion (F_8, 1027_ = 2.256, P = 0.022, η^2^_P_ = 0.017). Habitat matching was also influenced by interactions between trait diversity and ecoregion for FRic (FRic × ecoregion, F_8, 1015_ = 7.388, P < 0.001, η^2^_P_ = 0.055), FEve (FEve × ecoregion, F_8, 1027_ = 4.337, P < 0.001, η^2^_P_ = 0.033), and FDiv (FDiv × ecoregion, F_8, 1027_ = 2.322, P = 0.018, η^2^_P_ = 0.018). There was no evidence for a main effect or interaction of FDis for either environmental filtering or habitat matching (Appendix Tables S3 and S4). Ecoregion consistently had the strongest effect on environmental filtering and habitat matching, although effects were weaker for habitat matching (Appendix Tables S3 and S4).

### Disequilibrium-by-Traits

Functional trait predictors of environmental filtering and habitat matching varied by individual trait, trait category, and ecoregion (Figures 4 and 5). Environmental filtering was primarily influenced by habitat traits (7/9 ecoregions), with dispersal (4/9 ecoregions) and ecology (4/9 ecoregions) traits generally of secondary and tertiary influence, respectively (Figure 4). In contrast to environmental filtering, habitat matching was primarily influenced by both habitat (4/9 ecoregions) and dispersal (4/9 ecoregions) traits (Figure 5). Habitat traits were also commonly the secondary influence on habitat matching (5/9 ecoregions), with both dispersal (4/9 ecoregions) and ecology (4/9 ecoregions) traits of tertiary importance.

**Figure 4:**
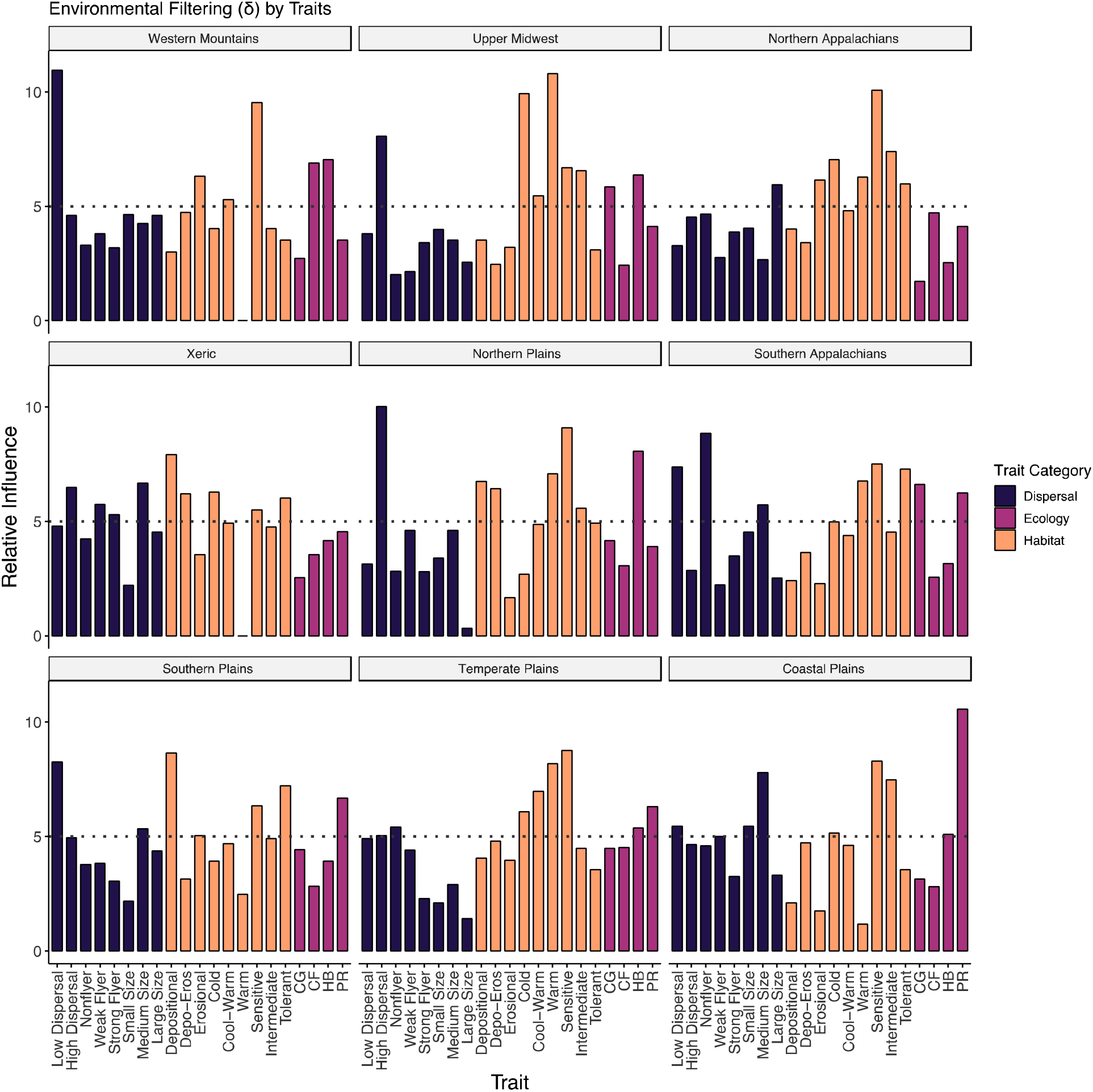
Relative influence of dispersal, habitat, and ecology functional traits for predicting environmental filtering (*δ*). The grey dashed line indicates a relative influence of 5.00. Select traits are abbreviated as: Depo-Eros = depositional-erosional, CG = collector-gatherer, CF = collector-filterer, HB = herbivore, and PR = predator. Ecoregion facet plots are arranged by approximate geographic position.

**Figure 5:**
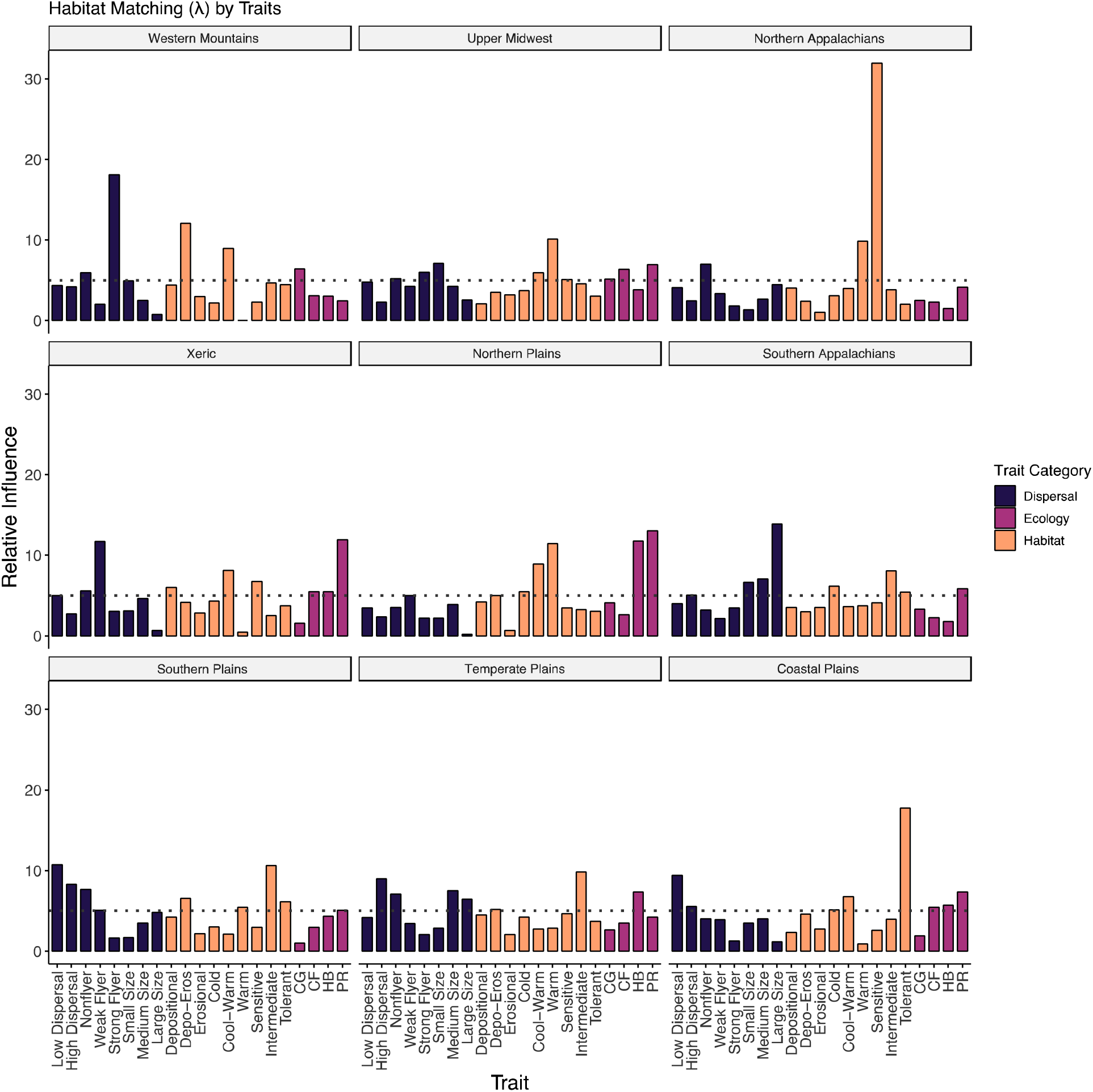
Relative influence of dispersal, habitat, and ecology functional traits for predicting habitat matching (*λ*). The grey dashed line indicates a relative influence of 5.00. Select traits are abbreviated as: Depo-Eros = depositional-erosional, CG = collector-gatherer, CF = collector-filterer, HB = herbivore, and PR = predator. Ecoregion facet plots are arranged by approximate geographic position.

### Traits-by-Environment

Predictors of functional traits varied by trait category and ecoregion (Figure 6). Dispersal traits were primarily influenced by network (39/72 trait-by-ecoregion combinations) and environmental (23/81 trait-by-ecoregion combinations) predictors, with environmental predictors also frequently of secondary influence (41/81 trait-by-ecoregion combinations) and landscape predictors of tertiary influence (39/81 trait-by-ecoregion combinations). Similarly, habitat traits were primarily influenced by network (48/81 trait-by-ecoregion combinations) and environmental (29/81 trait-by-ecoregion combinations) predictors; environmental predictors were commonly of secondary influence (37/81 trait-by-ecoregion combinations) and landscape predictors of tertiary influence (40/81 trait-by-ecoregion combinations). Ecology traits were primarily structured by environmental (17/36 trait-by-ecoregion combinations) and network (15/36 trait-by-ecoregion combinations) predictors. Each of environmental, landscape, and network predictors were frequently of secondary influence for ecology traits (environmental = 12/36, landscape = 10/36, network = 14/36 trait-by-ecoregion combinations), which was in contrast to dispersal and habitat traits where environmental predictors were most commonly of secondary influence; however, we again identified landscape predictors to most commonly be of tertiary importance (20/36 trait-by-ecoregion combinations).

**Figure 6:**
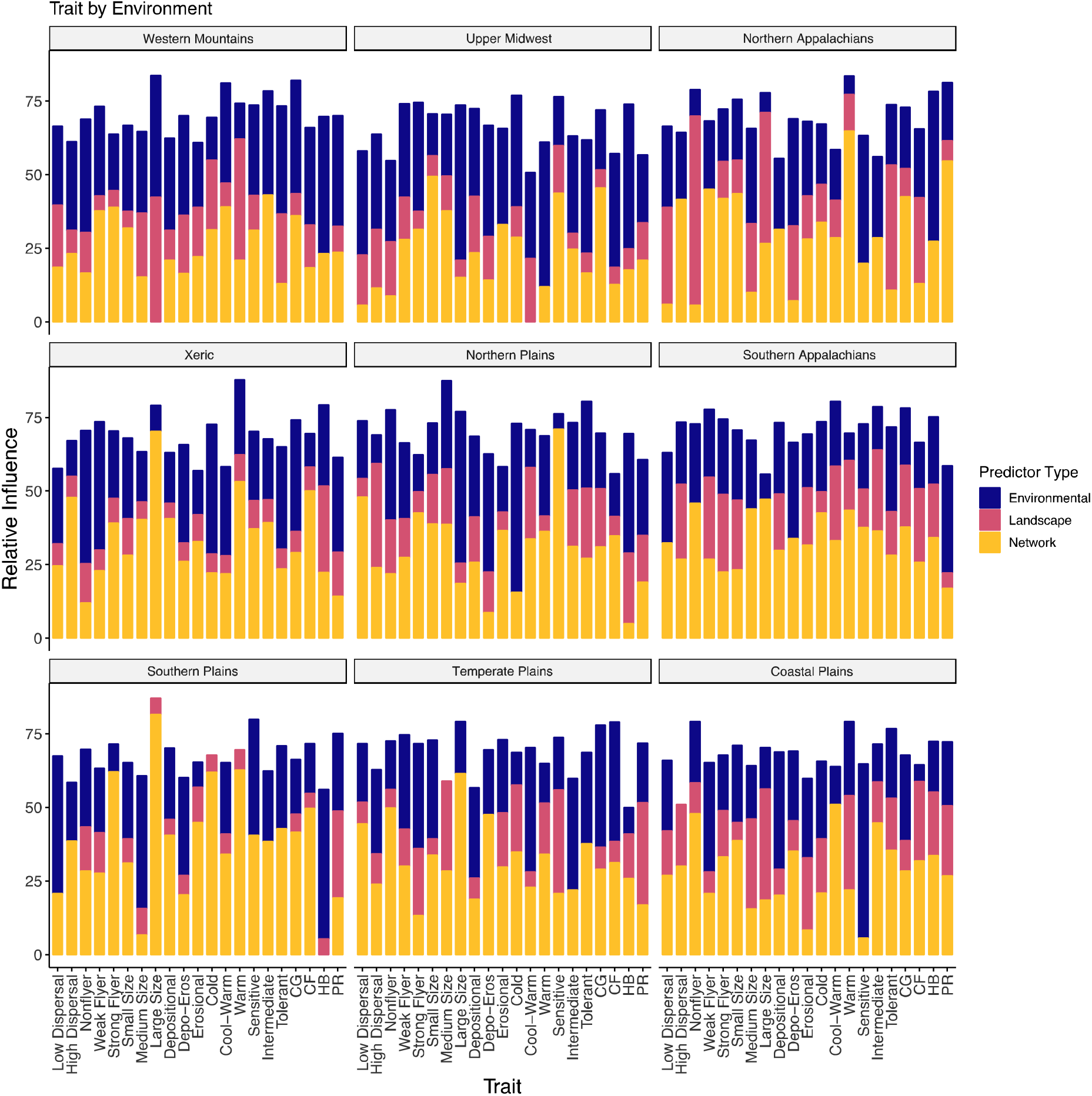
Summed relative influence of environmental, landscape, and network variables for predicting functional trait abundances. Sums equal the sum of all predictor variables within the respective category with a relative influence > 5.00.Traits are abbreviated as: Depo-Eros = depositional-erosional, CG = collector-gatherer, CF = collector-filterer, HB = herbivore, and PR = predator. Ecoregion facet plots are arranged by approximate geographic position.

## Discussion

Our study demonstrates spatial patterns of environmental filtering and habitat matching in communities across ecoregions at the macroecological scale, with functional traits providing a critical link between disequilibrium metrics and the environment. Support for this claim is derived from four key results. First, environmental filtering strongly varied by ecoregion, but habitat matching only exhibited weak spatial variation across ecoregions (**Q1**). Second, functional trait diversity also varied across ecoregion; however, patterns of functional trait diversity depended on the estimated metric (**Q2**). Third, we found that functional traits related to local habitat conditions were consistently the best predictor of environmental filtering across ecoregions, with dispersal traits also playing a role (**Q3**). Habitat matching was best predicted by both dispersal- and habitat-related traits (**Q3**). Additionally, functional trait diversity was primarily related to habitat matching and not environmental filtering (**Q3**). Finally, we found that the drivers of trait-environment relationships were context dependent, with predictors varying in identity and strength across ecoregions (**Q4**). With the available evidence, we suggest that environmental filtering and habitat matching depend on spatial context and associated changes and linkages with functional traits.

### Environmental Filtering

Both the identity and magnitude of functional trait predictors varied among ecoregions, which further suggests context dependency underlying the spatial patterns in environmental filtering and permissiveness. Context dependency is a common refrain within ecology (Leibold & Chase, 2017); however, there are explanations and paths forward through which context-dependency could be better understood. First, variation among ecoregions suggests a role of biogeography on regional and local communities, whereby landscape and environmental characteristics within an ecoregion affect the evolution of functional traits and community structure (Cavender-Bares et al., 2009; Cornell & Harrison, 2014; Leibold & Chase, 2017; Mittelbach & Schemske, 2015). Processes operating at biogeographical and regional scales could then eventually impact community responses at the local level; therefore, further investigation of the biogeography of these ecoregional species pools would provide more thorough background to explain functional trait evolution and resulting patterns in functional trait diversity (Cavender-Bares et al., 2009; Cornell & Harrison, 2014; Leibold & Chase, 2017; Mittelbach & Schemske, 2015). Second, given the link between organisms and ecology provided by functional traits (McGill et al., 2006; Spasojevic et al., 2014) and the utility for functional traits to better explain community assembly (Gianuca et al., 2018; Grönroos et al., 2013; Heino & Grönroos, 2014), focusing on habitat and dispersal traits should help us better understand why communities have certain functional traits as predictors. For example, communities in the northern plains and upper midwest ecoregions displayed strong signatures of environmental filtering and were both predicted by high dispersal and sensitive tolerance, but the northern plains were also predicted by warm water preference while the upper midwest was predicted by cold water preference (Figure 4). High dispersal and sensitive tolerance could suggest strong ecological selection (sensu Vellend 2010), resulting in species sorting despite potential mass effects through high dispersal abilities (Heino, Melo, et al., 2015; Leibold et al., 2004; Leibold & Chase, 2017; Shmida & Wilson, 1985). Lastly, combining trait and phylogenetic information can help elucidate the drivers of ecological patterns across space (Cavender-Bares et al., 2009; Gianuca et al., 2017, 2018; Leibold et al., 2010), although current limitations in taxonomic and spatial coverage for freshwater invertebrates limit this approach at macroecological scales. We used functional traits to help link patterns to potential ecological processes, but a functional perspective is not the only valid approach. Expanding open databases with phylogenetic data will help us to identify functional and phylogenetic relationships with environmental filtering, including the identification of concordant and discordant explanations provided by functional traits and community phylogenetics (e.g., Gianuca et al., 2018).

### Habitat Matching

Contrary to environmental filtering, there were no meaningful differences in habitat matching among ecoregions (Figure 5). All ecoregions displayed signatures overlapping both habitat matching (negative *λ*) and mismatch (positive Λ). These patterns were surprising, given mismatch is generally rare (Blonder et al., 2015); however, there are two points we would like to emphasize to contextualize our findings. First, the increased frequency of mismatch in our study could imply: (1) many communities have either undergone a perturbation pushing observed communities into mismatch or (2) communities have undergone or were undergoing disturbance events heterogeneously in space; both points are not necessarily mutually exclusive. A key limitation we recognize is that our current study did not and cannot evaluate temporal variation of the habitat matching and mismatch patterns or assess communities before and after disturbance events or regime shifts (Knight et al., 2020; Schriever et al., 2015; Walker et al., 2021). Such explicit considerations of community change through time and space would make characterizations of habitat (mis)matching more robust and should be investigated in future studies. Second, the lack of variation in habitat matching among ecoregions does not entail the lack of variation within ecoregions. Indeed, we found strongly matched and mismatched communities across the majority of ecoregions (7/9 ecoregions, Appendix Figure S5), with the exceptions (Southern Appalachians and upper midwest) still displaying overlap between habitat matching and mismatch (Appendix Figure S5). There was also no relationship between environmental filtering and habitat matching (Appendix Figure S5), suggesting independence between these two metrics. We did find frequent relationships between habitat matching and community-wide patterns of function trait diversity (Appendix Figures S3 and S4). When combined with habitat and ecology traits as influential predictors of habitat matching, this suggests that accounting for both the identity and distribution of functional traits is important for understanding the dynamics of habitat matching.

### Predictors of Functional Traits

We identified a link between functional traits and community disequilibrium, but the drivers of functional trait abundance depended on trait category and ecoregion. Dispersal traits were primarily influenced by network variables, which supports previous research documenting spatial influence on communities (Germain et al., 2017; Göthe et al., 2017; Grönroos et al., 2013; Heino & Grönroos, 2014; Horváth et al., 2019; Sarremejane et al., 2017). We found that mean annual flow, a measure of stream size and position within a dendritic network (Altermatt, 2013), was a common predictor of dispersal traits. Additionally, basin area, a measure of habitat amount (Fahrig, 2013) and incoming flows of nutrients and organisms (Altermatt, 2013; Heino, Melo, et al., 2015), was also a frequent driver of dispersal traits. It is important to emphasize that longitude and latitude were the most consistent drivers of dispersal traits, which suggests an important role for spatial position within the ecoregional metacommunity. These results suggest a dual influence of habitat amount and spatial configuration on dispersal traits, which follows previous work documenting hierarchical and interactive effects of multiple processes acting simultaneously on local communities (Astorga et al., 2011; Göthe et al., 2017; Heino et al., 2017; Murray Stoker & Murray Stoker, 2020). Forested land cover frequently influenced habitat traits, along with impervious surface cover. Both of these landscape factors can affect community assembly, diversity, and composition, though often with contrasting ecological consequences. Forested land cover is generally related to increased diversity (Budnick et al., 2019; Wahl et al., 2013), and our results suggest that forested land cover was linked to rheophilic, thermal, and pollution tolerance traits. Similarly, impervious surface cover typically reduces diversity and alters community composition (Barnum et al., 2017; Gianuca et al., 2018; Walker et al., 2021), and we found rheophilic and thermal traits were predicted by impervious surface cover.

### Biogeography, Species Pools, and Functional Traits

Regional species pools have long been recognized as important mediators of local responses (Astorga et al., 2011; Cornell & Harrison, 2014; Heino et al., 2017; Leibold & Chase, 2017; Mittelbach & Schemske, 2015), and that was reflected in our study. We found that community composition (Appendix Figure S6) and functional trait diversity varied by ecoregion, with some ecoregions overlapping in composition and others were comparatively distinct (Appendix Figure S6), confirming established differences in ecoregional metacommunity structure (Murray Stoker & Murray Stoker, 2020); however, we were unable to incorporate several facets of the structure and diversity of regional species pools. We evaluated regional species pools in terms of community composition and functional diversity, but evolutionary history and phylogenetic relationships can be an important component of regional species pools (Cavender-Bares et al., 2009; Cornell & Harrison, 2014; Leibold & Chase, 2017; Mittelbach & Schemske, 2015). We also evaluated function trait diversity through categorical trait assignments, as there were limitations in the available data; however, intraspecific trait variation and quantitative assessment of functional traits could better define functional trait structure of the regional pool and provide a more accurate link between organisms to ecology (Des Roches et al., 2018; Leibold & Chase, 2017). We also defined our species pools based on ecoregions, which were defined based on biotic and abiotic characteristics (USEPA 2016a). Studies frequently set the basin as the regional species pool (Altermatt et al., 2013; Grönroos et al., 2013; Heino & Grönroos, 2014), but this delineation would have been impractical in our study: multiple sites were only sampled in a fraction of the basins (< 10%), precluding a robust evaluation and assessment. Differences in the definition of regional species pools notwithstanding, we contend our methodology was valid for answering our research questions within the restrictions of the study design. More broadly, it is possible that variation in the regional species pools underlies the context dependency of environmental filtering and habitat matching. For example, expansions and retractions of species ranges driven by glaciations can structure taxonomic and genetic diversity (Hickerson et al., 2010), resulting in regions varying in species endemism and diversity (e.g., Hamilton & Morse, 1990) and potentially constraining trait evolution. Further investigation of regional species pools and associated functional structure, particularly the explicit inclusion of evolutionary history and intraspecific trait variation, could help explain this ostensible context dependency.

### Limitations

We recommend that inferences and predictions are considered within the appropriate limitations of our study. Like any macroecology study, there is an inherent trade-off between spatial and temporal coverage. While we were able to capture a substantial spatial scale, we were not able to evaluate temporal changes in this study. Species interactions can also have an important effect on community assembly and disequilibrium (Blonder et al., 2015; Wisz et al., 2013). While species interactions are theoretically incorporated into habitat matching calculations (Blonder et al., 2015), a more explicit consideration and assessment would be useful. Functional traits can be used for a more mechanistic link in community ecology (McGill et al., 2006; Spasojevic et al., 2014), yet there was consistent context dependency in community Disequilibrium-by-Traits and Trait-by-Environment linkages. Rather than approaching context dependency as a problem, we suggest embracing it. Given the pervasive and inherent variation in the drivers of community disequilibrium and functional trait abundances, understanding how and why these ecological phenomena are context dependent is necessary. We also want to acknowledge that our study is largely descriptive, but we contend that descriptive studies at the macroecological scale are necessary to set the frame of reference and guide further investigations into currently unanswered questions.

### Conclusion

Our study demonstrates the patterns and drivers of environmental filtering and habitat matching on a macroecological scale. We found that both environmental filtering and permissiveness were common signatures at the ecoregional scale. In contrast, habitat matching did not vary across ecoregions and many communities could be pushed to either equilibrium or disequilibrium. We also found that functional trait diversity varied among ecoregions, with effects on habitat matching but not environmental filtering. Linkages between community disequilibrium and functional traits were identified, along with the concomitant identification of environmental variables that predicted functional trait structure. In summary, our study demonstrates the contingency of environmental filtering and habitat matching and the underlying trait predictors and trait-environment relationships. Given the expanding and strengthening threats to freshwater biodiversity with global change (Booth et al., 2016; Cid et al., 2022; Dudgeon, 2019; Dudgeon et al., 2006), it is imperative to identify vulnerable communities and potential causal relationships operating at local and regional scales. We aim for this work to provide the foundation on which trait-environment relationships can be further quantified and causal explanations established in the context of community disequilibrium and applied to the conservation and management of freshwater systems.

## Acknowledgements

Our research would not have been possible without the extensive efforts and expertise of the dedicated field crews, laboratory staff, data management and quality control staff, analysts and many others from the EPA, states, tribes, federal agencies, universities, and other organizations that conducted the National Rivers and Streams Assessment (2008-2009). We are not affiliated with the USEPA, and the views and opinions expressed in this paper are solely of the authors and do not reflect any official position of the agency. We wish to recognize that the rivers and streams sampled for this study are a part of landscapes that form the ancestral homes of many distinct Native groups. We encourage other researchers to recognize the traditional stewards of their local environment in what is now the United States at https://native-land.ca/.

## Biosketches & Author Contributions

<u>David Murray-Stoker</u> is a PhD candidate at the University of Toronto with interests in community and evolutionary ecology, pedagogy, and the history and philosophy of science.

<u>Kelly Murray-Stoker</u> is a PhD candidate at the University of Toronto interested in community ecology, taxonomy and systematics of caddisflies, and decolonial and feminist ecologies.

<u>Fan Peng Kong</u> is a post-bachelor researcher from the University of Toronto Mississauga currently working as an energy coach and client representative at an energy efficiency non-profit.

<u>Fathima Amanat</u> is currently completing an ecology and evolution specialist and minor in environmental management and has always been curious about the interconnectedness of nature. She is keen to continuously learn about it in her personal and academic life.

## Author Contributions

DMS designed the project and led the data analysis and writing. DMS, KMS, FPK, and FA interpreted results, and KMS, FPK, and FA contributed to manuscript writing and revisions.

## Data Availability

All data, metadata, and code are deposited on Zenodo (Murray-Stoker et al. 2022).

