## Appendix S1 for "Environmental filtering and habitat (mis)matching of riverine invertebrate metacommunities"

**Supplemental Methods**

*Sampling design and methods*

We used data from the 2008-2009 National Rivers and Streams Assessment (NRSA) conducted by the United States Environmental Protection Agency. Sites for the NRSA were selected by implementing a generalized random tessellation stratified design, which gives each potential site a known probability of selection and inclusion in the study (USEPA 2016a). Moreover, the design was explicitly structured to incorporate the range of variation for rivers and streams, and site selection and inclusion further controlled to ensure sites were distributed among different Strahler orders (USEPA 2016a). In total, 1924 sites were selected for the NRSA and distributed across nine ecoregions: coastal plains, northern Appalachians, northern plains, southern Appalachians, southern plains, temperate plains, upper midwest, western mountains, and xeric; ecoregions for the NRSA were assigned following the classification system proposed by Omernik (1987). Data were collected for benthic macroinvertebrates, water chemistry, and physical habitat characteristics. Land cover attributes for the basin of each site were derived from the National Hydrography Dataset Plus (USEPA 2016a, b).

Sites were sampled following standardized protocols between late May-early October of 2008 and 2009 (USEPA 2016a). Sampling reach lengths were proportional to the wetted-width of the channel. For wadeable rivers and streams, the sampling reach was 40 times the wetted-width of the channel, and streams with a wetted-width less than 3.5 m had a sampling reach of 150 m (USEPA 2016c). For larger non-wadeable rivers and streams, sampling reach lengths were also set to 40 times the wetted-width of the channel and up to a maximum length of 2000 m (USEPA 2016c). For each site, the center of the sampling reach (i.e., X-site) was marked and 11 equally-spaced transects were demarcated along the sampling reach. Water chemistry samples were collected at the X-site at a depth of 0.5 m, with physical habitat and substrate characteristics estimated visually for each of the 11 transects and then averaged. Macroinvertebrate samples were collected in composite, whereby individual samples from each of the 11 transects were combined into the composite sample (USEPA 2016c). For wadeable rivers and streams, macroinvertebrates were collected by sampling a 0.09 m^2^ quadrat for 30 s using a D-frame kick net (500 μm mesh), with the position of the quadrat selected at random (USEPA 2016c). Similarly, macroinvertebrates were collected from non-wadeable rivers and streams using a single 1-m sweep of the dominant habitat type in the transect with a D-frame kick net (500 μm mesh, USEPA 2016c). All composite macroinvertebrate samples were preserved in 95% ethanol and identified (primarily to genus, USEPA 2016c). Sampling for the NRSA required 85 crews, with all crews required to attend a training course before sampling. Additionally, 10% of all sites were resampled for quality control (USEPA 2016a).

*Niche variables*

We used air T_max_ annual, air T_min_ annual, pH, and conductivity to estimate species niches in the disequilibrium analysis (Blonder et al., 2015). Temperature can have profound effects on macroinvertebrate physiology and ecology, including thermoregulation, metabolism, activity, development, and thermal tolerance levels (Gullan & Cranston, 2014; Merritt et al., 2019). We included pH and conductivity as these variables broadly related to pollution and disturbance (Dodds & Whiles, 2010; Merritt et al., 2019; Paul & Meyer, 2001; Walsh et al., 2005). While there is not as direct a link between pH or conductivity to macroinvertebrate physiology as there is for the temperature variables, we used pH and conductivity because they can be related to a broad suite of environmental factors and ecological disturbances (e.g., urbanization, pollution, sedimentation, eutrophication, hypoxia/anoxia; Dodds & Whiles, 2010; Gullan & Cranston, 2014; Merritt et al., 2019; Paul & Meyer, 2001; Walsh et al., 2005). These environmental factors and ecological disturbances, in turn, can affect macroinvertebrate physiology and ecology, which ultimately affects community assembly and concomitant species distributions. Including pH and conductivity allowed us to therefore capture a wide range of influences on macroinvertebrates using as few variables as possible. Because the disequilibrium analysis is sensitive to the number of variables used to define species’ niches, we followed the recommendation by Blonder et al. (2015) to include as few and ecologically- and physiologically-relevant variables as possible; our choice of variables captured a wide range of environmental influences on macroinvertebrate physiology and ecology.

*Caveats:* Although having water temperature at annual timescales would have been helpful, air temperature is still a useful measure for macroinvertebrate physiology. For example, many macroinvertebrates have both aquatic and terrestrial life stages and temperature would still have an effect on their niche and distribution (Gullan & Cranston, 2014; Merritt et al., 2019). Water temperature data is logistically challenging to collect at large spatial and temporal scales, like in our study, so air temperature is used by default as it is the only variable that can be collected or calculated at macroecological scales, spatial and/or temporal.

*Note on variable selection:* While exchanging variables could change the multidimensional niche space and alter estimates of environmental filtering and habitat matching, those estimates would be for different components of the multidimensional niche. Our study was focused on understanding community assembly and responses to environmental change in river and stream ecosystems, and we contend that the variables we selected capture the relevant axes within the multidimensional niche for the purposes of our study. We would suggest that studies include variables that are relevant to the axes of the multidimensional niche being examined, as no single study can reliably incorporate the full multidimensional niche without distorting the niche space used by the disequilibrium pipeline to quantify environmental filtering and habitat matching.

*Environmental, landscape, and network variables*

We selected environmental, landscape, and network variables to identify influential drivers of DisEQ-by-trait and trait-by-environment relationships. Environmental variables were selected as measures of local habitat conditions. Total nitrogen and total phosphorus are frequently collected (Bini et al., 2014; Gianuca et al., 2017, 2018; Heino et al., 2003, 2017; Walker et al., 2021) and can be used to estimate nutrient enrichment or even indicate eutrophication of a river or stream (Dudgeon, 2019; Paul & Meyer, 2001; Vörösmarty et al., 2010; Walsh et al., 2005); total nitrogen and phosphorous levels are commonly elevated with increasing anthropogenic disturbance (Paul & Meyer, 2001; Walsh et al., 2005). Dissolved organic carbon was selected because it can be an important resource within the food web in rivers and streams (Dodds & Whiles, 2010). We used natural, algal, and macrophyte cover as they can serve as both habitat and an energy source (Dodds & Whiles, 2010; Walker et al., 2021), with large woody debris included primarily as a source of habitat (Dodds & Whiles, 2010; Walker et al., 2021).

Landscape variables were selected as measures of land cover characteristics within the basin of each site. Forested land cover is tightly linked to habitat quality in rivers and streams, with a positive relationship between forested land cover and habitat quality (Allan, 2004; Dodds & Whiles, 2010); there is a notable exception between forested land cover and desert streams, where reduced or even the absence of forested land cover is expected even in high-quality habitats (Allan, 2004; Dodds & Whiles, 2010). Agricultural land cover can relate to elevated nutrient inputs and eutrophication of rivers and stream ecosystems (Allan, 2004; Dodds & Whiles, 2010), where agricultural land cover is negatively related to habitat quality and biodiversity. Both urban and impervious surface cover were selected to estimate the effects of urbanization and broader anthropogenic disturbance (Allan, 2004; Dodds & Whiles, 2010; Paul & Meyer, 2001; Walsh et al., 2005); while both variables are generally related, there can be a discordance between urban and impervious surface cover (i.e., not all impervious surface cover is classified as urban; Allan, 2004; Dodds & Whiles, 2010; Paul & Meyer, 2001; Walsh et al., 2005). Urban and impervious surface cover typically reduce habitat quality and biodiversity.

Network variables were selected to identify site position and connectedness within the metacommunity and regional pool. Latitude and longitude relate to the spatial location of each site. Basin area is the total area that drains into the site and can estimate incoming flows of organisms and nutrients from tributaries. Mean annual flow can be a useful surrogate for position in the dendritic networks often displayed by streams and rivers (Altermatt, 2013). Actual measurements of stream flow can also be more reliable estimates of stream size and position instead of more coarse designations (e.g., Strahler order; Altermatt, 2013). The mean and range of elevation within the basin are related to site isolation and barriers to dispersal within the network (Kärnä et al., 2015; Tonkin et al., 2018). Site centrality is the mean distance of a site to all other sites within the network, providing a measure of site connectedness. While more commonly applied to sites within the same basin (e.g., Altermatt et al., 2013), site centrality can be useful for evaluating communities at larger spatial scales (e.g., Murray‐Stoker & Murray‐Stoker, 2020). Site centrality was calculated as the mean geodesic distance on an ellipsoid using the `geosphere` package (Hijmans, 2019). Larger values for site centrality indicate longer dispersal distances and lower connectedness while smaller values indicate shorter dispersal distances and higher connectedness.

*Descriptions of functional trait diversity metrics*

We measured functional trait diversity for each community using four indices: (1) functional richness; (2) functional divergence; (3) functional evenness; and (4) functional dispersion. Functional richness (FRic) is the portion of trait space occupied by the community (Schleuter et al., 2010; Villéger et al., 2008). Functional evenness (FEve) measures the regularity of the distribution of traits within occupied trait space using a minimum spanning tree (Schleuter et al., 2010; Villéger et al., 2008). Functional divergence (FDiv) is the proportion of trait space occupied by extreme trait values (Schleuter et al., 2010; Villéger et al., 2008). Functional dispersion (FDis) is the weighted mean distance of individual taxa in the community to the centroid of all species in multidimensional trait space, and simultaneously measures trait dissimilarity and evenness within the community (Botta‐Dukát, 2005; Laliberté & Legendre, 2010). FDis is weighted by species abundances, shifting the centroid towards species that are more abundant. Communities with high FDis are composed of evenly distributed, dissimilar traits while communities with low FDis are composed of unevenly distributed, similar traits.

**Table S1:** Seven trait categories and associated trait modalities used for evaluating functional trait diversity and assessing functional trait abundances. Traits were compiled from Poff et al. (2006).

| **Trait Category** | | **Modality** |
| --- | --- | --- |
| Dispersal | Dispersal | (1) Low (< 1 km flight/travel before oviposition) |
|  |  | (2) High (> 1 km flight/travel before oviposition) |
|  | Flying Strength | (1) Nonflyer |
|  |  | (2) Weak flyer |
|  |  | (3) Strong flyer |
|  | Body Size | (1) Small |
|  |  | (2) Medium |
|  |  | (3) Large |
| Habitat | Rheophily | (1) Depositional |
|  |  | (2) Depositional-erosional |
|  |  | (3) Erosional |
|  | Thermal Preference | (1) Cold |
|  |  | (2) Cool-warm |
|  |  | (3) Warm |
|  | Pollution Tolerance | (1) Sensitive |
|  |  | (2) Intermediate |
|  |  | (3) Tolerant |
| Ecology | Functional Feeding Group | (1) Collector-gatherer (including shredders*) |
|  |  | (2) Collector-filterer |
|  |  | (3) Herbivore |
|  |  | (4) Predator |

*Only one shredder was retained in our dataset (*Lepidostoma*) and comprised less than 0.50% of total abundance.

**Table S2:** Summary statistics [mean (1 standard deviation)] for predictor variables used in the boosted regression tree (BRT) analyses. Ecoregions are abbreviated as: coastal plains = CPL, Northern Appalachians = NAP, northern plains = NPL, Southern Appalachians = SAP, southern plains = SPL, temperate plains = TPL, western mountains = WMT, upper midwest = UMW, and xeric = XER. We selected seven environmental variables [total nitrogen (μg L^-1^, Total N), total phosphorus (μg L^-1^, Total P), dissolved organic carbon (mg L^-1^, DOC), large woody debris (m^3^ per 100 m, LWD), natural streambed cover (proportion, NAT Cover), algal cover (proportion, ALG Cover), and macrophyte cover (proportion, AQM Cover)], four landscape variables [forested cover (% of basin area, % For), agricultural cover (% of basin area, % Ag), urban cover (% of basin area, % Urb), and impervious surface cover (% of basin area, % ISC)], and seven network [latitude, longitude, basin area (km^2^), mean annual flow (m^3^ s^-1^, Mean Flow), mean basin elevation (m, Mean Elevation), range of basin elevation (m, Range Elevation), and site centrality (distance to centroid in km, Centrality)] variables.

| **Variable** | **Ecoregion** | | | | | | | | |
| --- | --- | --- | --- | --- | --- | --- | --- | --- | --- |
|  | **CPL** | **NAP** | **NPL** | **SAP** | **SPL** | **TPL** | **UMW** | **WMT** | **XER** |
| **Environmental** | | | | | | | | | |
| Total N | 1283.29 (1810.43) | 664.01  (838.80) | 1314.80 (4167.53) | 1046.70 (2309.52) | 1809.89 (3422.72) | 3558.77 (4082.09) | 1558.52 (1922.09) | 251.54  (601.52) | 782.36 (2297.26) |
| Total P | 199.55  (336.10) | 56.99  (114.48) | 230.96  (584.58) | 87.24  (195.46) | 455.05 (1280.51) | 238.53  (495.88) | 150.54  (381.49) | 126.42  (911.44) | 166.96  (542.55) |
| DOC | 10.08 (14.70) | 4.02 (2.91) | 7.85 (5.21) | 2.37 (1.82) | 4.48 (2.73) | 5.57 (8.58) | 9..41 (10.11) | 1.93 (1.90) | 2.94 (2.37) |
| LWD | 2.61 (5.18) | 5.37 (8.24) | 1.13 (4.15) | 4.82 (13.11) | 2.83 (10.98) | 3.96 (10.08) | 3.52 (5.77) | 6.30 (13.96) | 1.15 (2.50) |
| NAT Cover | 0.39 (0.31) | 0.54 (0.34) | 0.21 (0.24) | 0.37 (0.21) | 0.24 (0.22) | 0.23 (0.18) | 0.29 (0.19) | 0.52 (0.40) | 0.40 (0.37) |
| ALG Cover | 0.04 (0.12) | 0.04 (0.09) | 0.05 (0.13) | 0.01 (0.04) | 0.06 (0.13) | 0.04 (0.10) | 0.04 (0.10) | 0.04 (0.09) | 0.07 (0.14) |
| AQM Cover | 0.10 (0.17) | 0.05 (0.09) | 0.11 (0.18) | 0.03 (0.07) | 0.07 (0.16) | 0.05 (0.16) | 0.07 (0.14) | 0.08 (0.14) | 0.12 (0.18) |
| **Landscape** | | | | | | | | | |
| % For | 39.64 (25.47) | 65.69 (27.18) | 8.76 (15.91) | 62.23 (25.61) | 12.42 (16.59) | 9.78 (15.66) | 38.20 (24.42) | 65.86 (21.74) | 31.29 (26.64) |
| % Ag | 24.69 (26.66) | 15.48 (22.07) | 18.60 (21.58) | 20.34 (21.66) | 26.56 (25.55) | 72.16 (24.71) | 31.68 (29.80) | 0.34 (1.07) | 1.78 (5.17) |
| % Urb | 11.65 (19.86) | 9.29 (15.47) | 1.72 (1.45) | 11.45 (18.96) | 4.41 (6.37) | 8.08 (12.88) | 7.07 (14.59) | 1.37 (2.53) | 1.59 (3.56) |
| % ISC | 2.80 (7.35) | 2.12 (4.43) | 0.29 (0.17) | 2.31 (5.54) | 0.83 (2.75) | 1.55 (3.80) | 1.52 (5.86) | 0.24 (0.49) | 0.42 (0.95) |
| **Network** | | | | | | | | | |
| Latitude | 33.71 (3.37) | 42.79 (3.37) | 45.45 (1.63) | 36.80 (2.10) | 37.21 (3.39) | 41.35 (2.79) | 44.91 (1.71) | 41.71 (4.34) | 39.42 (3.25) |
| Longitude | −86.09 (6.84) | −74.88 (3.84) | −103.84 (3.30) | −83.95 (5.53) | −99.71 (2.59) | −92.30 (4.37) | −89.63 (3.25) | −114.35 (6.27) | −114.09 (4.42) |
| Basin Area | 74.44  (260.34) | 79.45  (226.01) | 4423.64 (7566.12) | 171.52  (638.19) | 15335.91 (28135.59) | 525.25 (1715.87) | 243.81  (567.93) | 688.62 (1755.52) | 5671.01 (20665.27) |
| Mean Flow | 0.83 (2.63) | 1.23 (3.31) | 1.57 (3.74) | 2.20 (8.37) | 18.62 (50.78) | 0.85 (1.50) | 1.30 (2.29) | 1.88 (4.73) | 7.12 (18.97) |
| Mean Elevation | 71.42  (46.25) | 373.20  (178.08) | 1089.50 (451.04) | 358.49  (232.92) | 996.35  (655.02) | 319.56  (95.01) | 336.65  (80.00) | 1965.92 (859.14) | 1957.40 (559.95) |
| Range Elevation | 42.84  (39.73) | 285.52  (261.51) | 665.34  (585.38) | 237.58  (246.51) | 1058.55 (1271.65) | 85.99  (71.80) | 94.89  (66.57) | 982.02  (595.57) | 1383.13 (728.23) |
| Centrality | 626.61 (208.16) | 566.39 (196.10) | 456.08 (160.82) | 601.24 (209.48) | 579.67 (214.03) | 650.70 (227.91) | 727.24 (217.67) | 620.51 (227.77) | 683.80 (225.49) |

**Table S3:** Summary of the ANCOVAs comparing environmental filtering against functional trait diversity [functional richness (FRic), functional evenness (FEve), functional divergence (FDiv) and functional dispersion (FDis)]. We report the degrees of freedom (df), F-statistics calculated from type III sums-of-squares, and the effect size (partial eta-squared, η^2^_P_) for each term in the ANCOVAs.

| **Term** | **df** | **F** | **P-value** | **η^2^_P_** |
| --- | --- | --- | --- | --- |
| **FRic** | | | | |
| FRic | 1 | 0.075 | 0.784 | 0.000 |
| Ecoregion | 8 | 77.853 | < 0.001 | 0.380 |
| FRic × Ecoregion | 8 | 1.591 | 0.123 | 0.012 |
| Residuals | 1015 |  |  |  |
| **FEve** | | | | |
| FEve | 1 | 0.256 | 0.613 | 0.000 |
| Ecoregion | 8 | 82.861 | < 0.001 | 0.392 |
| FEve × Ecoregion | 8 | 0.544 | 0.823 | 0.004 |
| Residuals | 1027 |  |  |  |
| **FDiv** | | | | |
| FDiv | 1 | 0.076 | 0.783 | 0.000 |
| Ecoregion | 8 | 77.745 | < 0.001 | 0.377 |
| FDiv × Ecoregion | 8 | 2.256 | 0.022 | 0.017 |
| Residuals | 1027 |  |  |  |
| **FDis** | | | | |
| FDis | 1 | 0.001 | 0.979 | 0.000 |
| Ecoregion | 8 | 93.792 | < 0.001 | 0.418 |
| FDis × Ecoregion | 8 | 0.244 | 0.982 | 0.002 |
| Residuals | 1046 |  |  |  |

**Table S4:** Summary of the ANCOVAs comparing habitat matching against functional trait diversity [functional richness (FRic), functional evenness (FEve), functional divergence (FDiv) and functional dispersion (FDis)]. We report the degrees of freedom (df), F-statistics calculated from type III sums-of-squares, and the effect size (partial eta-squared, η^2^_P_) for each term in the ANCOVAs.

| **Term** | **df** | **F** | **P-value** | **η^2^_P_** |
| --- | --- | --- | --- | --- |
| **FRic** | | | | |
| FRic | 1 | 2.373 | 0.124 | 0.002 |
| Ecoregion | 8 | 7.915 | < 0.001 | 0.059 |
| FRic × Ecoregion | 8 | 7.388 | < 0.001 | 0.055 |
| Residuals | 1015 |  |  |  |
| **FEve** | | | | |
| FEve | 1 | 4.277 | 0.039 | 0.004 |
| Ecoregion | 8 | 3.276 | 0.001 | 0.025 |
| FEve × Ecoregion | 8 | 4.337 | < 0.001 | 0.033 |
| Residuals | 1027 |  |  |  |
| **FDiv** | | | | |
| FDiv | 1 | 6.092 | 0.014 | 0.006 |
| Ecoregion | 8 | 1.802 | 0.073 | 0.014 |
| FDiv × Ecoregion | 8 | 2.322 | 0.018 | 0.018 |
| Residuals | 1027 |  |  |  |
| **FDis** | | | | |
| FDis | 1 | 0.968 | 0.325 | 0.001 |
| Ecoregion | 8 | 0.669 | 0.719 | 0.005 |
| FDis × Ecoregion | 8 | 0.695 | 0.696 | 0.005 |
| Residuals | 1046 |  |  |  |

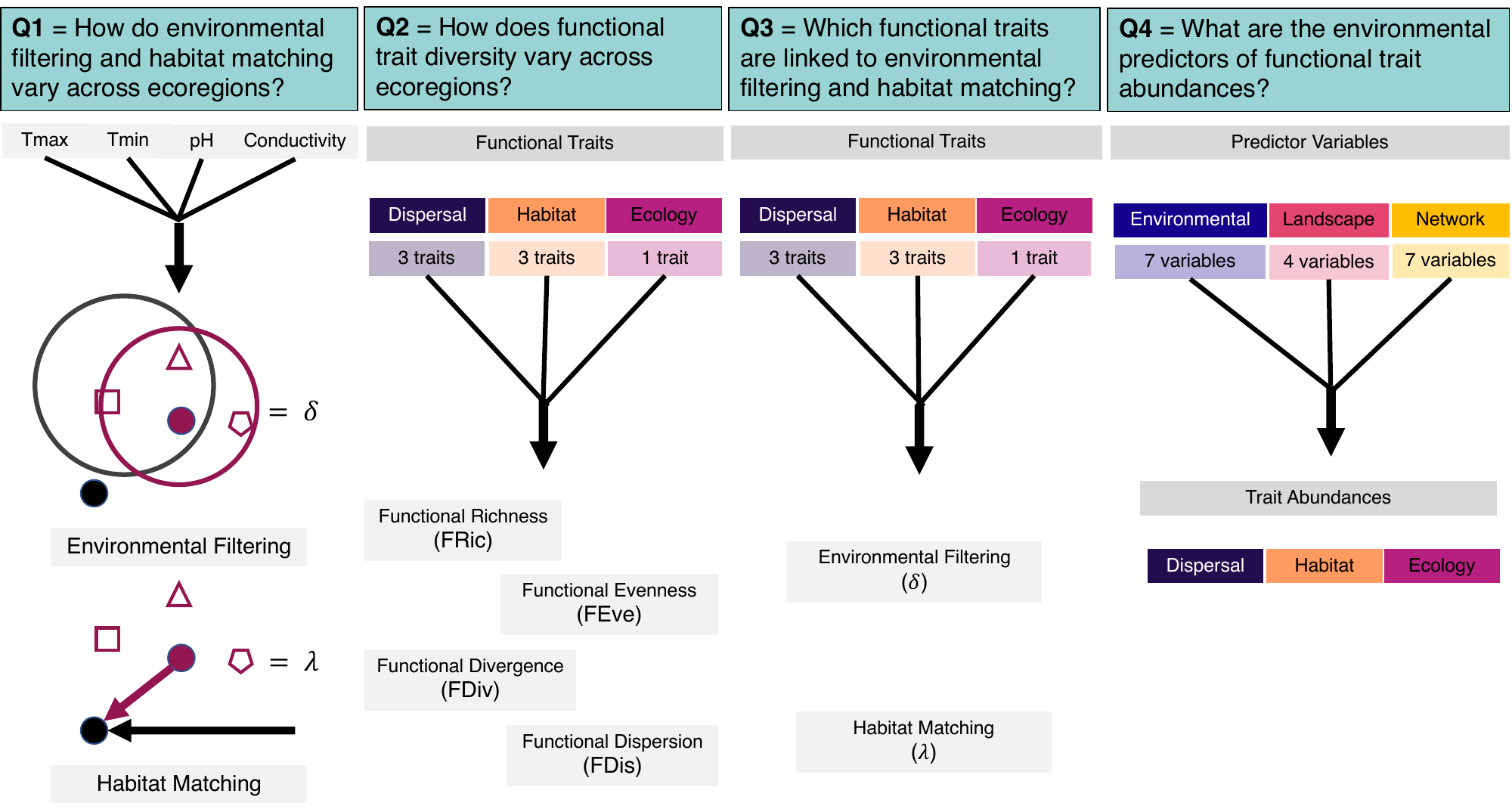

**Figure S1:** Conceptual diagram of the data and workflow to answer each of the four research questions for this study. (Q1) We used T_max_ annual, T_min_ annual, pH, and conductivity to estimate species’ niches and then calculate environmental filtering (𝛿) and habitat matching (𝜆). Observed community niche volume (Δ) is measured as the niche space occupied by the community and accounts for each species’ niche breadth (red circle) and the centroid for the community is inferred (red dot) from the positions of observed community members (red square, triangle, and pentagon) within the niche space (Blonder et al., 2015). The community niche volume and centroid of null assemblages (niche volume = black circle, centroid = black dot) are calculated by resampling communities of equal richness from the regional pool (Blonder et al., 2015), scaling the observed niche volumes by the null expectations. As the observed community (red circle) has a smaller niche volume (𝛿 < 0) than the null expectation (gray circle), the observed community was structured by environmental filtering (Blonder et al., 2015). A larger niche volume (𝛿 > 0) than the null expectation would indicate environmental permissiveness (Blonder et al., 2015). Similarly, observed community habitat matching (Λ) is the vector between the centroid (red dot) and the null centroid (black dot), with the observed vector indicated by the red arrow and a null vector indicated by the black arrow. After scaling the observed vectors by the null vectors, the observed community would be classified as matching the habitat (i.e., habitat matching) because the observed vector is shorter than the null vector (𝜆 < 0; Blonder et al., 2015). In contrast, habitat mismatch would be indicated if the observed vector was longer than the null vector (𝜆 > 0; Blonder et al., 2015). Measures of environmental filtering and habitat matching were calculated for each community, with comparisons made among ecoregions. (Q2) We used three functional traits with a total of 21 trait modalities to quantify functional trait diversity for each community and then compared functional trait diversity among ecoregions. (Q3) Using the same functional traits from Q2, we calculated the abundance (i.e., count) of each trait in each community, and we then identified functional traits that were predictors of environmental filtering and habitat matching (taken from Q1) across ecoregions. (Q4) Using selected environmental, landscape, and network variables, we identified predictors of functional trait abundances (from Q3) across ecoregions.

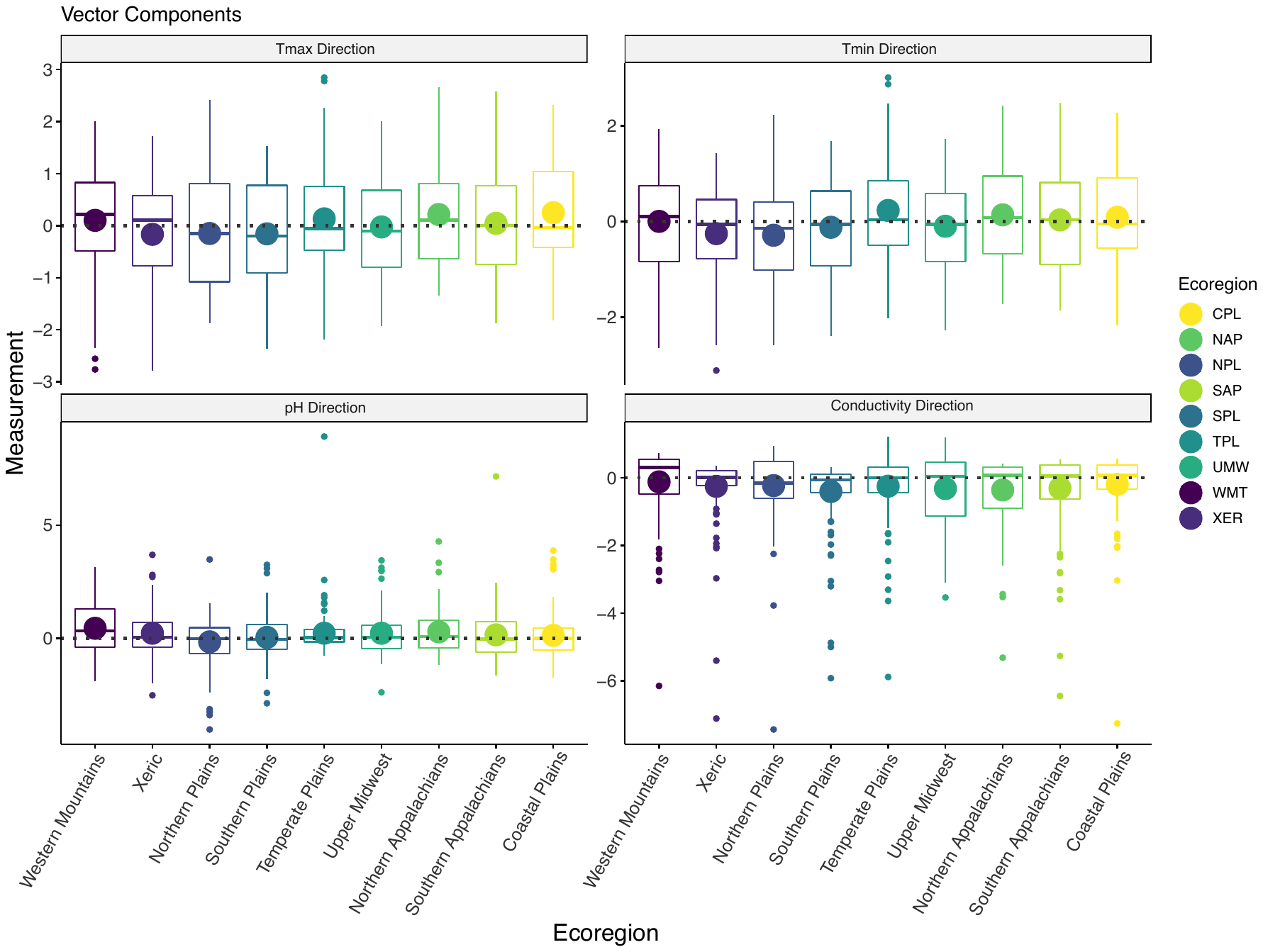

**Figure S2:** Facet plot of habitat matching vector components by ecoregion (Tmax = annual maximum air temperature, T_min_ = annual minimum air temperature, pH, and conductivity). Large circles represent the mean and boxplots display the interquartile range (0.25, 0.75), minimum, and maximum, with smaller circles indicating outliers. Negative values indicate habitat matching and positive values indicate habitat mismatch. Ecoregions are arranged on the x-axis in approximate position based on longitude.

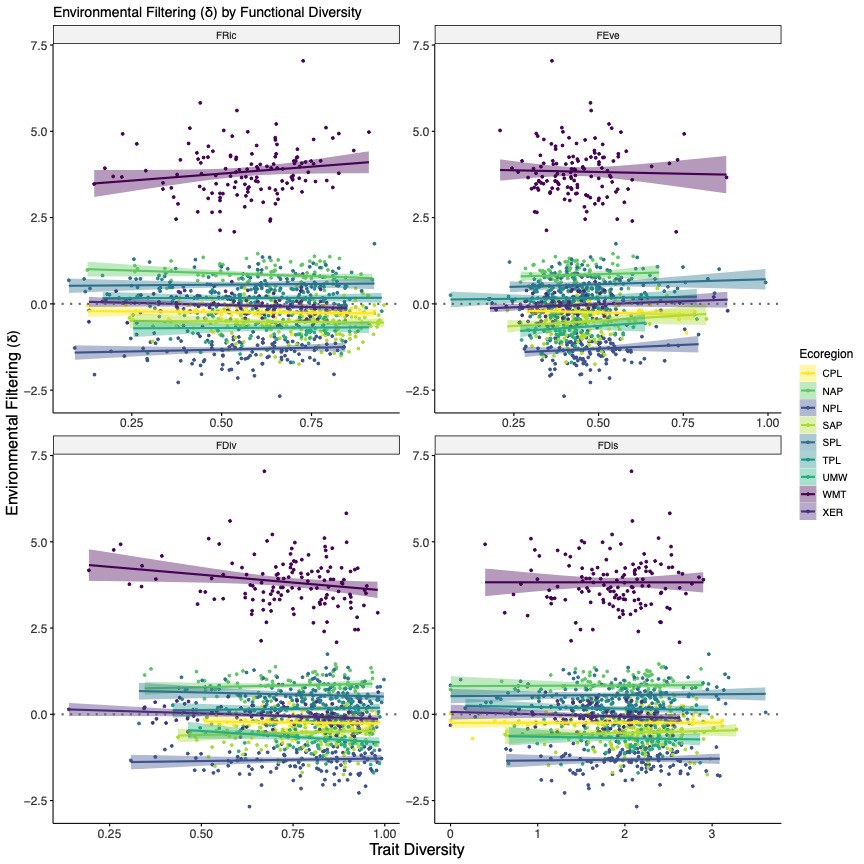

**Figure S3:** Facet plot of environmental filtering against functional richness (FRic), functional evenness (FEve), functional divergence (FDiv) and functional dispersion (FDis), with separate lines for each ecoregion. Lines are lines-of-best fit (± 95% confidence interval) from ANCOVA models, allowing the slope and intercept to vary by ecoregion. Negative values indicate environmental filtering and positive values indicate environmental permissiveness. Model statistics are provided in Appendix Table S3. Ecoregions are abbreviated in the legend as: coastal plains = CPL, Northern Appalachians = NAP, northern plains = NPL, Southern Appalachians = SAP, southern plains = SPL, temperate plains = TPL, upper Midwest = UMW, western mountains = WMT, and xeric = XER.

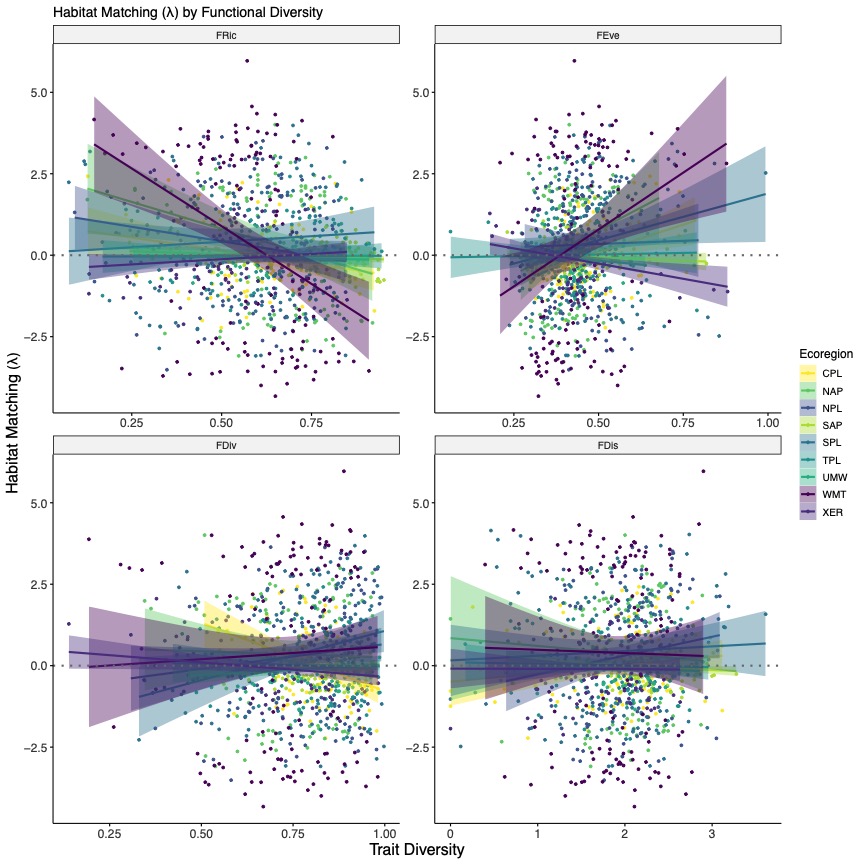

**Figure S4:** Facet plot of habitat matching against functional richness (FRic), functional evenness (FEve), functional divergence (FDiv) and functional dispersion (FDis), with separate lines for each ecoregion. Lines are lines-of-best fit (± 95% confidence interval) from ANCOVA models, allowing the slope and intercept to vary by ecoregion. Negative values indicate environmental filtering and positive values indicate environmental permissiveness. Model statistics are provided in Appendix Table S4. Ecoregions are abbreviated in the legend as: coastal plains = CPL, Northern Appalachians = NAP, northern plains = NPL, Southern Appalachians = SAP, southern plains = SPL, temperate plains = TPL, upper Midwest = UMW, western mountains = WMT, and xeric = XER.

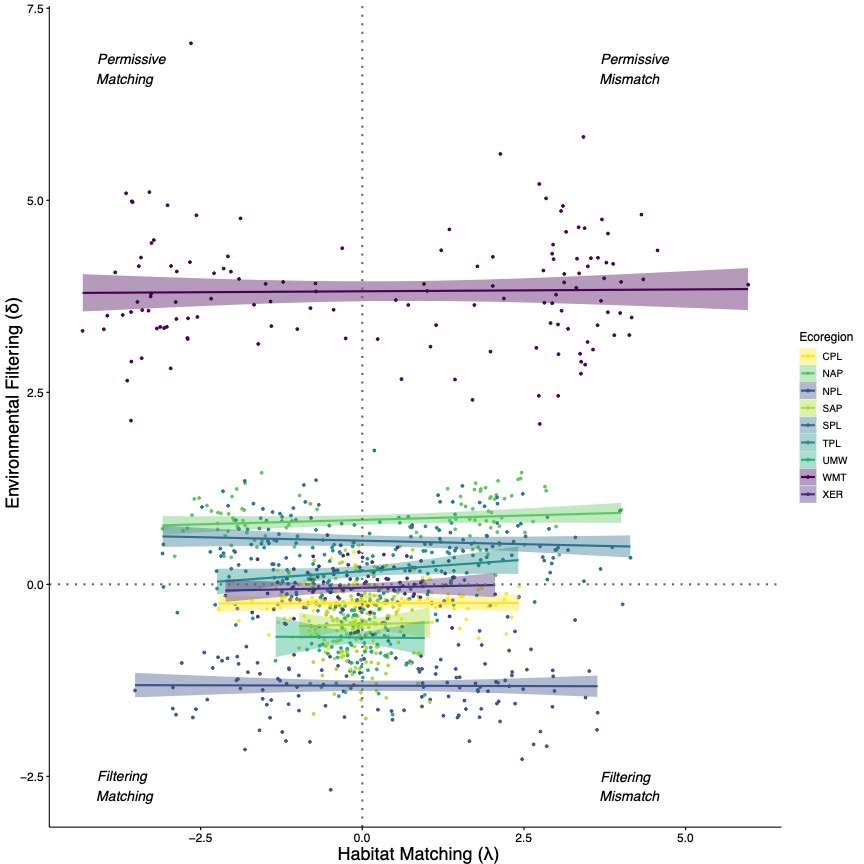

**Figure S5:** Plot of environmental filtering against habitat matching, with separate lines for each ecoregion. Lines are lines-of-best fit (± 95% confidence interval) from a linear mixed-effects model, whereby environmental filtering was regressed against habitat matching and ecoregion was fitted as a random intercept; we allowed slopes to vary in the figure to aid in interpretation. There was no relationship between environmental filtering and habitat matching (𝛽 = 0.005, standard error = 0.008, df = 1068.026, t = 0.554), with ecoregion explaining the majority of the variation (R^2^ ≅ 0.90). We fitted the linear mixed-effects model using `lmer()` in the `lme4` (Bates et al., 2015) and `lmerTest` (Kuznetsova et al., 2017) packages, with model assumptions evaluated using `check_model()` in the `performance` package (Lüdecke et al., 2021). Ecoregions are abbreviated in the legend as: coastal plains = CPL, Northern Appalachians = NAP, northern plains = NPL, Southern Appalachians = SAP, southern plains = SPL, temperate plains = TPL, upper Midwest = UMW, western mountains = WMT, and xeric = XER.

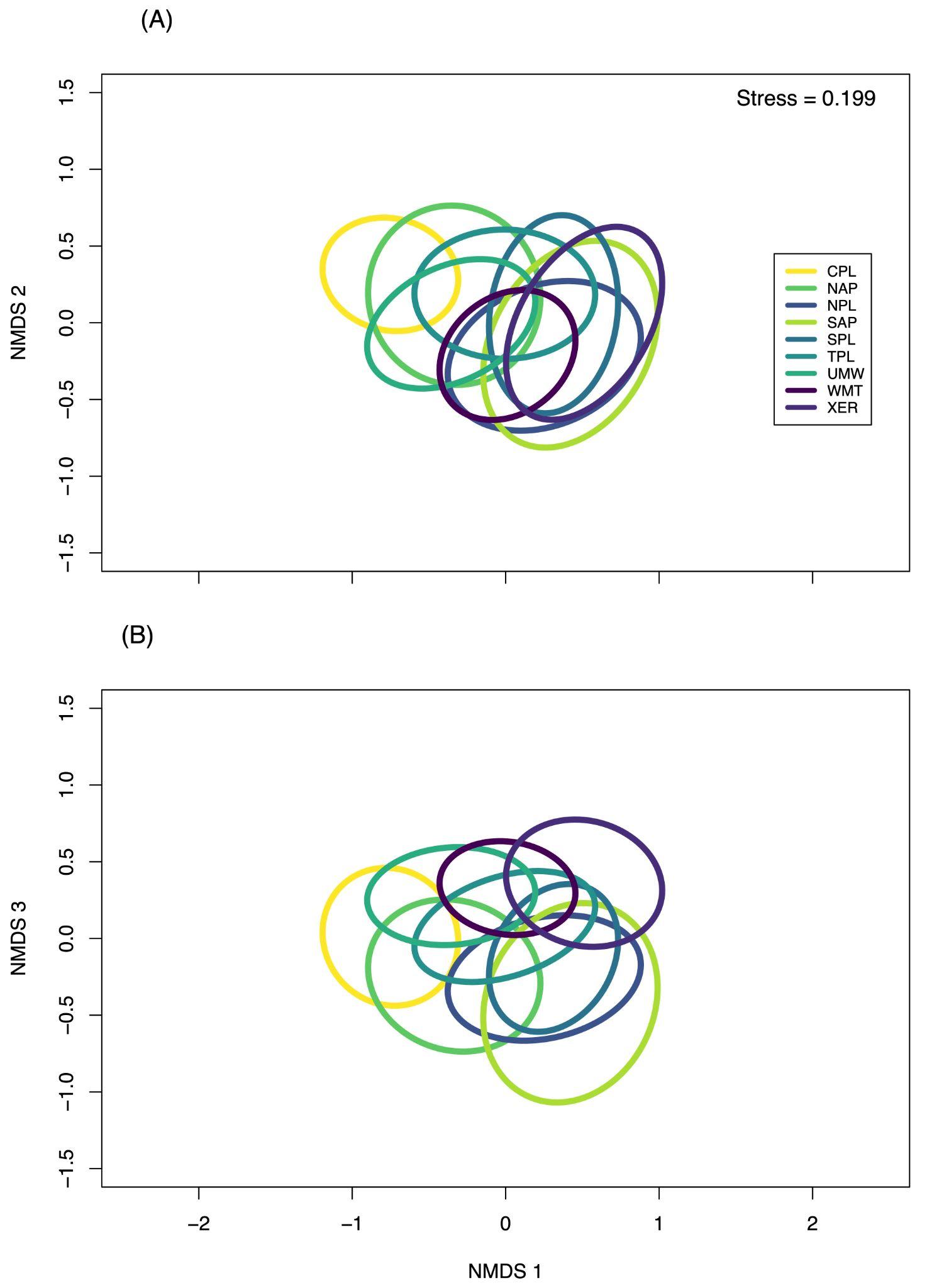

**Figure S6:** Non-metric multi-dimensional scaling plot of the Bray-Curtis dissimilarity of macroinvertebrate community composition among ecoregions. Three dimensions were required to achieve adequate stress in ordination space, and dimensions 1 and 2 are presented in panel (A) and dimensions 1 and 3 are presented in panel (B); dimension 1 is held constant on the x-axis for reference. Each ecoregion displayed a distinct community composition. (PERMANOVA, R^2^ = 0.083, P < 0.001). We conducted the PERMANOVA with 10000 permutations using the `adonis()` function (Oksanen et al., 2020). Ecoregions are abbreviated in the legend as: coastal plains = CPL, Northern Appalachians = NAP, northern plains = NPL, Southern Appalachians = SAP, southern plains = SPL, temperate plains = TPL, upper Midwest = UMW, western mountains = WMT, and xeric = XER.
